# Sumatran orangutan mothers differ in the extent and trajectory of their expression of maternal behaviour

**DOI:** 10.1101/2024.11.25.625129

**Authors:** T Revathe, Roger Mundry, Sri Suci Utami-Atmoko, Tazkia Umaira Aprilla, Maria A. van Noordwijk, Marlen Fröhlich, Paul-Christian Bürkner, Caroline Schuppli

## Abstract

Mothers play a crucial role in the early development and survival of mammalian offspring, and differences in maternal care may differentially affect offspring’s development. Whereas previous research has primarily focused on biological and socioecological factors to understand population-level variation in maternal behaviour, the individual as a source of variation remains understudied. We investigated between-individual variation in the average expression of, and plasticity in, six maternal behaviours in Sumatran orangutans, using 15 years of behavioural data. We found that mothers differed substantially in the average expression of four maternal behaviours, even after controlling for socioecological conditions, biological state characteristics, and the offspring’s influence on these behaviours. Furthermore, not controlling for these confounding effects exaggerated or masked between-individual variation. Mothers also substantially differed in how they adjusted three of the maternal behaviours during offspring development, meaning that mothers differed in behavioural plasticity. Our results suggest that Sumatran orangutan mothers are constrained in the average expression of maternal behaviours and their plastic responses, potentially resulting in consistent differences among mothers, otherwise called maternal personality. Our findings highlight that individual variation around the population mean in maternal behaviour is more than noise and presents opportunities to study novel evolutionary processes that shape maternal behaviour.

## 1. Introduction

In all mammals, mothers are crucial for offspring’s healthy development, survival into adulthood, and, eventually, successful reproduction [1–5]. Mothers achieve this by nursing, grooming, defending, and provisioning food to their offspring, through imparting knowledge on their offspring, and providing coalitionary and reproductive support to their offspring [6–11]. Ultimately, maternal behaviour is an adaptive trait affecting the fitness of both the mother and the offspring [9].

Maternal behaviour towards offspring is particularly varied in long-lived species – such as primates [7] – in which mothers support their offspring through a lengthy period of development, and the mother-offspring relationship often extends far beyond weaning [9, 11]. In several primate species, the extent of expression of a maternal behaviour does not always completely overlap among all individuals in a population, i.e., there may be behavioural variation among mothers [12] (henceforth, between-individual variation in the average expression of a maternal behaviour). Though such between-individual variation among mothers can arise from the differences in mother-offspring characteristics and the immediate socioecological conditions under which they are observed, previous research suggests that there may be fundamental differences among mothers, which may remain consistent across different conditions, including across one’s different offspring and different social and ecological environments [13,14].

Consistent differences among females in their maternal behaviour can emerge through differences in social and environmental conditions they faced during their own early developmental period or can be intrinsic [15,16]. In humans, consistent behavioural differences among individuals are often termed personality, and this term is applied more and more to nonhuman species as well [17–20]. Although maternal behaviours are rarely included in the traits used to assign personality, they are interesting candidates to investigate consistent behavioural differences. Differences among mothers may lead to differences in the quantity and quality of offspring-directed maternal behaviours. In human and non-human primates, the quantity and quality of offspring-directed maternal behaviours may influence offspring’s pace of development, exploration tendencies, cognitive abilities, learning, skill acquisition, social behaviour, and ultimately survival [21–25], (but see [26]). Therefore, beyond the immediate presence of the mother [3–5], the nature of her investment plays an important role in the successful rearing of her offspring.

Consistent differences between individuals in maternal behaviour over time does not mean that mothers will not exhibit short-term variation in their behaviour within their limits. Indeed, maternal behaviour is usually flexible in nature, in that individuals alter the expression of their behaviour in response to changes in their ecological and social environment [12]. Maternal behavioural expression is, therefore, seldom constant, even within one offspring’s developmental period. Flexibility in the expression of maternal behaviour is expected to be especially pronounced in long-lived, large-brained (relative to body size) species, as they are more likely to face fluctuating socio-ecological conditions during their lifetime and have a greater cognitive potential to exhibit behavioural flexibility [27–29]. Long-lived, large-brained species were thus hypothesised to achieve adaptation to prevailing conditions through being behaviourally flexible (otherwise called behavioural plasticity) [27]. Primate mothers, for example, flexibly vary the expression of maternal behaviours in response to experienced ecological conditions such as food availability or predation risk [30,31] and social conditions such as social unit size, dominance structure, and presence of kin or a male in the social unit [32–34]. Primate mothers also vary their maternal behaviours as a function of their biological and social characteristics such as body condition, parity, age, and dominance status [35–37], and of their offspring’s characteristics – most importantly, offspring age [35,37–40]. In other words, often there is within-individual variation or behavioural plasticity in maternal behaviour when measured over the ranges of different predictors.

Behavioural plasticity in maternal behaviour is hypothesized to positively affect female lifetime reproductive success [6]. Two recent studies found that not all primate mothers exhibit the same degree of plasticity in maternal behaviours, such as offspring-directed communication, time spent in proximity to their offspring, retrieving and restraining their offspring, and mother leaving their offspring [26,41]. Though behavioural plasticity and between-individual variation in maternal behavioural plasticity remain largely understudied in primates, a study on at least one non-primate species has revealed that females who exhibited higher phenotypic behavioural plasticity were also found to have a higher reproductive success [42].

Though behavioural traits from an individual’s perspective can be viewed through the above two contrasting ideas of consistency and flexibility, most behaviours are neither rigidly fixed nor unlimitedly plastic, i.e., they are shaped by both the flexible and consistent nature of individuals [12]. Estimating between-individual variation in the average expression of a behaviour is often complicated by the fact that between-individual variation is confounded by the factors influencing within-individual variation (i.e., confounding effects of maternal behaviour; see below). Therefore, only through the simultaneous investigation of between- and within-individual variation can we fully understand the role of individuals in the expression of maternal behaviour.

To understand the adaptive value of between- and within-individual variation, it is important to investigate these in wild animals. However, estimating between- and within-individual variation in maternal behaviour using field data is challenging, as data on wild individuals are rarely complete due to the practical challenges associated with uniformly sampling the individuals through an extended period of offspring development, such as in primates. For example, individuals cannot always be found in the study area or only a limited number of individuals can be observed simultaneously due to logistic constraints. This introduces sampling discrepancies among individuals, resulting in a significant risk of wrongly inferring the presence of between-individual variation in maternal behaviour when there is none (i.e., false-positive difference); or a failure to detect the presence of between-individual variation when it is actually present (i.e., false-negative similarity), further discussed in [26,43]. These constraints raise the question of whether studies actually measure consistent, between-individual variation when they do not control for the confounding effects (i.e., offspring age, offspring sex, mother’s parity, sample size, etc., as discussed above).

The behavioural reaction norm approach [42,44–47] offers a way to reliably estimate between- and within-individual variation in maternal behaviour simultaneously, especially when faced with sampling discrepancies and/or confounding effects. The approach entails controlling for confounding effects and estimating between- and within-individual behavioural variation – without isolating them, using a mixed model approach (Supplementary Material 1, Box 1), which allows one to estimate between-individual variation relative to within-individual variation. The approach has been successfully applied to several behavioural traits, including maternal behaviours, movement behaviour, calving date, egg laying date, chick provisioning behaviour and male territorial aggressive behaviour, in different species [42,47–52]. Studies on mammalian maternal behavioural variation using behavioural reaction norms are sparse, (but see [26, 41]). As long-term data on individually identified animals and large sample sizes are necessary to identify individual variation, such studies are challenging in species with slow life-histories and prolonged maternal care.

In this study, we aimed to shed light on the extent to which maternal behaviour in wild Sumatran orangutans (*Pongo abelii*) is shaped by individual variation by integrating both consistent individual differences and variation in individuals’ plasticity. We investigated between-individual variation in the average behavioural expression and between-individual variation in plasticity in six maternal behaviours while controlling for previously established predictors of, as well as the offspring’s potential influence on, maternal behaviour, using longitudinal and cross-sectional behavioural data. Previous studies on individual variation in maternal behaviour in primates did not control for all known confounding effects before quantifying individual variation [13,26,39,41,53,54]. Therefore, one of the aims of this study was to identify whether not controlling for known confounding effects (i.e., the biological characteristics of the mother and/ or offspring, the prevailing socioecological conditions experienced by the mother-offspring pair, sample size, and sampling scheme; see Methods) would influence the estimation of between-individual variation in the average expression of maternal behaviour.

Sumatran orangutans are large-brained and long-lived arboreal apes, living up to at least 50-60 years in the wild [55,56]. Sumatran orangutan females start reproducing from around 15 years of age; with a duration of about 8 years, their inter-birth interval is the longest among all the non-human primates [57]. Sumatran orangutans are an excellent study system to investigate individual maternal behavioural variation, as mothers provide care to each of their offspring for about 6-9 years without help from other individuals [58], through extensive caretaking behaviours such as carrying, bridging, nest-sharing, and maintaining proximity to offspring, while serving as role models for the offspring’s acquisition of learning-intense subsistence skills such as feeding and nest building [59–62].

In line with the hypothesis that individuals may be constrained in their behavioural expression due to their genetic and/or developmental background [15,16], a previous study on wild and captive orangutans found that orangutans show between-individual variation in communicative behaviours, including offspring-directed maternal communication and responsiveness [41]. Based on this result and the results from studies on other primates [26,32,39,63],

➢ we predicted that Sumatran orangutan mothers will differ in the average expression of maternal behaviours even after controlling for known confounding effects. Since a mother may face differential demands from different offspring, we predicted that a part – but not all – of the variation among mothers in their expression of maternal behaviour will be explained by offspring’s identity.

In a previous work on the study population [40], we found that offspring’s biological characteristics (i.e., offspring age, offspring sex) and experienced socioecological factors (i.e., food availability, association size, and presence of a male) had significant effects on several maternal behaviours. We thus hypothesised that the estimation of between-individual variation in the average expression of maternal behaviour is influenced by whether these known confounding effects are controlled for in the analysis.

➢ We predicted that the estimated between-individual variation in the average expression of maternal behaviour will be exaggerated or, alternatively, masked and will have a higher uncertainty associated with it when confounding effects are not controlled for than when they are appropriately controlled for in the analysis.

The extent of behavioural plasticity is hypothesised to be positively correlated with brain size [27]. Accordingly, orangutans have been found to show considerable plasticity in maternal behaviours and maternal gestural communication [40,41]. Since behavioural plasticity is expected to balance the costs and benefits of maternal behaviour, and as it is unlikely that all mothers face similar pressures during the long offspring developmental period, it is likely that the plastic responses are different among Sumatran orangutan mothers. In fact, a recent study found that mothers differed in how they altered their responsiveness towards offspring depending on the context of interaction [41].

➢ We predicted that Sumatran orangutan mothers will differ in their plasticity in maternal behaviours – i.e., mothers will differ in their maternal behavioural trajectories over offspring age. Since the differences among offspring in their demands from the mother may vary during the developmental period, we predicted that offspring identity will contribute to maternal behavioural plasticity but not fully account for it.

To carry out this study, we focused on maternal behaviours related to proximity maintenance, skill acquisition, and locomotory support [40] in 15 wild Sumatran orangutans at the Suaq Balimbing monitoring station in South Aceh, Indonesia.

## 2. Methods

### (a) Study site and data collection

To study individual variation in maternal behaviour, we used long-term data on six maternal behaviours, namely *initiation* and *termination of body contact* and *close proximity*, *carrying*, and *feeding in close proximity* (described in Supplementary Material 1, Table S1), collected on wild Sumatran orangutan mother-offspring pairs (*Pongo abelii*). The data were collected between 2007 and 2022 in the Suaq Balimbing research area, which is a coastal peat swamp forest in the Gunung Leuser National Park in South Aceh, Indonesia. These behaviours are commonly expressed by primate mothers and are used to measure the dynamics of the mother-offspring relationship in many primate and non-primate species [26,40,53,64–67].

Behavioural data were collected during focal follows, conducted continuously from morning nest to night nest when possible, based on an established protocol for orangutan data collection (https://www.ab.mpg.de/571325/standarddatacollectionrules_suaq_detailed_jan204.pdf).

During the focal follows, scan sampling [68] was conducted every two minutes during which the activity of the focal individual (i.e., a mother with a dependent offspring) and distances (measured in classes of 0 m (contact), >0-2 m, 2-5 m, 5-10 m, and 10-50 m) between the focal individual and all the association partners (of any age) were noted down. A party was defined as two or more individuals who were within 50 m of one another. When the distance class changed between two scans, the observers noted down the identity of the individual initiating the change. Further details about the study site and data collection are provided in previous publications [58,61,69].

### (b) Sample size

We had behavioural data on 15 mothers, for nine of whom we had data on one dependent offspring (between birth and eight years of age), and for six others we had data on more than one offspring (three mothers with offspring of either sex; 3 mothers with either male or female offspring; Supplementary Material 2, Table S1). Though data on multiple offspring per mother are ideal for detecting differences among mothers that are independent of offspring characteristics (e.g., offspring sex), such datasets are difficult to obtain in species with long interbirth intervals. Including data on multiple offspring for some mothers and data on only a single offspring for others can conflate between-mother differences and between-offspring differences, which can make it harder to tease apart their effects. To avoid erroneously over or underestimating differences between the mothers, we conducted two sets of analyses: in the first set of analyses, we included data on all the mother-offspring pairs (i.e., data on some mothers with multiple offspring while others with only one offspring); in the second set of analyses, we retained data on only those mothers with multiple offspring (i.e., data on all mothers with multiple offspring). The latter set of analyses was conducted only for those behaviours for which we had data on multiple offspring per mother for at least five mothers to get reliable random effects estimates of mother and offspring identities.

We operationalised maternal behaviour as the proportion of scans in which the mother showed a behaviour, where the denominator controls for the opportunities available to the mother to show that behaviour (for e.g., number of scans during which a mother moved was the denominator for the behaviour *carrying*; Supplementary Material 1, Table S1). As a result of opportunistic sampling and as we only included follows during which the mother had the opportunity to show the analysed maternal behaviour at least once, the number of focal follows and total observation duration differed among the mothers and across behaviours (Supplementary Material 2, Table S1, Figure S1). The number of focal follows per behaviour ranged between 622 and 868 (i.e., the number of days during which focal individuals were followed, and the behaviour could be assessed), which equalled between 5853.7 *h* and 7915.1 *h of* observation time (Supplementary Material 2, Figure S1). The range of offspring age during which maternal behaviour could be recorded (i.e., sampling scheme) varied substantially among the offspring (Supplementary Material 2, Figure S2).

### (c) Statistical analysis

#### Estimating between-individual variation in average maternal behavioural expression and plasticity

We used a generalized linear mixed effects model (GLMM) to quantify between-individual variation in the average expression of a behaviour and between-individual variation in behavioural plasticity. In a recent study on the Suaq orangutan population, we found that certain offspring characteristics (offspring age [continuous], offspring sex [binary; male/female] and socioecological factors (prevailing food availability [henceforth, FAI; continuous], average association size during a follow [continuous], and male presence during a follow [binary; presence/absence]) were significant predictors of different maternal behaviours, but not mother’s parity [40]. Therefore, as detailed in the introduction, any variation among mothers in their maternal behaviour is likely to be at least partly caused by the variation introduced by these predictors. Hence, we modelled each maternal behaviour as a function of the respective known significant predictors of that behaviour. Further details about these predictors are available in [40; Supplementary Material 3, Table S1]. The sampling scheme discrepancies in our study (Supplementary Material 2, Figure S2) additionally underscore the importance of including offspring age as a predictor [43]. We z-standardized the continuous predictors to a mean of zero and a standard deviation of one, and we dummy-coded and mean-centered offspring sex and male presence/absence to ease model convergence and interpretation [70; see footnote of Supplementary Material 4, Table S1]. We included a random intercept of mother identity and a nested random intercept of offspring identity within mother identity, as there were mothers with more than one offspring (see above). The estimated standard deviation of random intercepts of mother identity gives us the extent of between-individual variation in the average expression of maternal behaviour. The estimated standard deviation of random intercepts of offspring identity gives us the extent of variation among offspring of the same mother. A non-zero estimate of the standard deviation of random intercepts of the mother and offspring identities would mean that offspring of the same mother differentially influence maternal behaviour but also that mothers behave in a consistent manner towards each of their offspring. A non-zero estimate for the mother but not for the offspring identity would mean that mothers behave in a consistent manner towards each of their offspring and that offspring of the same mother do not differentially influence maternal behaviour. Alternatively, a non-zero estimate for the offspring but not for the mother identity would mean that offspring of the same mother differentially influence maternal behaviour and that mothers do not behave in a consistent manner with each of their offspring. We further included random slopes of offspring age within mother identity and offspring identity. The estimated standard deviation of random slopes with mother identity gives us the extent of between-individual variation in maternal behavioural plasticity. We could not calculate repeatability (calculated as variance among individuals/variance within individuals over time; [50,71]) to get population-level estimates of the fraction of maternal behavioural variation that is due to between-individual variation, as models included complex random effect structure with multiple random slopes.

#### Modelling procedure

We fitted the GLMMs explained above (henceforth, full model) using a Bayesian framework in R, version 4.2.2 [72], using Stan, version 2.32.5 [73,74], via the brms package [75], which retrieves posterior distributions of estimated parameters. We fitted the models using a beta-binomial error distribution for *initiation* and *termination of body contact* and *close proximity* (events), and with a beta error distribution for *carrying* and *feeding in close proximity* (which are both states rather than events), both with a logit link function. We included the numbers of scans in which the mother showed a behaviour as the number of success and the opportunities to show a behaviour (Supplementary Material 1, Table S1) as the number of trials in the beta-binomial models. We operationalized the dependent variables as proportions in the beta models. To examine the goodness-of-fit of the models, we computed Bayesian *R*^2^ [76] using the *bayes_R2*function.

#### Statistical support for random effects

In addition to each of the full models, we fitted a model excluding the random intercepts of mother identity and offspring identity nested within mother identity and random slope of offspring age (henceforth, fixed effects model) and one including the random intercepts of mother identity and offspring identity nested within mother identity but excluding the random slope of offspring age (henceforth, random intercept model). We then compared these models using the leave-one-out cross-validation (LOO-CV) [77] as implemented in the loo package (version 2.7.0) using the *loo* function. LOO-CV allowed us to examine whether the out-of-sample predictive performance of a model is improved by the addition of model terms (here, the random intercepts and slopes), which is analogous to the Akaike information criterion [78]. We chose the more parsimonious model whose Δ*ELPD* ± 2 × *SE* did not overlap zero to draw inferences about between-individual differences in maternal behaviour.

#### Influence of confounding effects

To understand whether and how not controlling for known confounding effects affect the estimation of between-individual variation in the average expression of maternal behaviour, we fitted another model to each of the maternal behaviours without any fixed effects (i.e., not controlling for the known confounding effects) but with the random intercepts of mother identity and offspring identity nested within mother identity. We then compared the estimated between-individual variation in the average expression of maternal behaviour and the uncertainty associated with it between the full model and the above model. We assessed these results via visual inspection.

We report the population-level effects in the supplementary material (Supplementary material 4, Table S1, Figure S1).

## 3. Results

### Between-individual variation in the average expression of maternal behaviour

After having controlled for mother-offspring characteristics and socioecological conditions as well as accounting for the offspring’s influence on maternal behaviour, we found that in four of the maternal behaviours, namely *contact termination*, *close proximity termination*, *carrying*, and *feeding in close proximity*, there was significant and consistent (across one’s offspring) between-individual variation among mothers in the expression of these behaviours(Supplementary Material 5, Table S1) and the posterior distributions of standard deviation of the random intercept of mother identity being away from zero (Supplementary Material 6, Figure S1). Between-individual standard deviations for mothers ranged between 0.32 and 0.9 across these models (Table 1). Offspring of the same mother contributed to, but did not fully explain, the variation in mothers’ expression of maternal behaviours (SD of the random intercept of offspring ranged between 0.47 and 0.71, Table 1, Figure 1, Supplementary Material 6, Figure S1). Furthermore, mothers differed in the average expression of maternal behaviour across behaviours, and there was a significant (negative) correlation between the random effect intercepts of mother identity in the *carrying* model and the model for *feeding in close proximity* (but not between any of the random effect intercepts of any of the other behaviours; Supplementary Material 7, Table S1).

**Figure 1.**
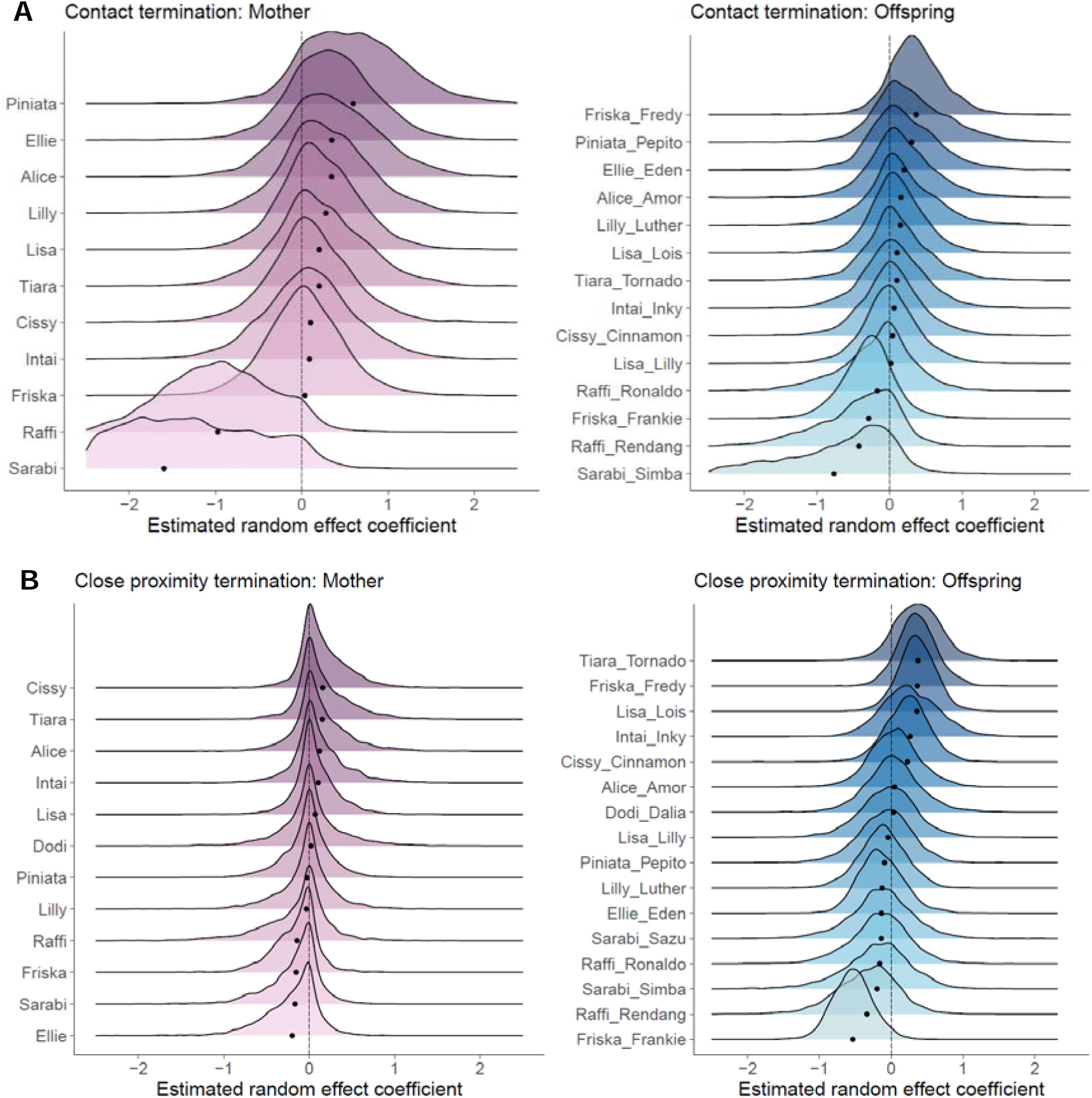
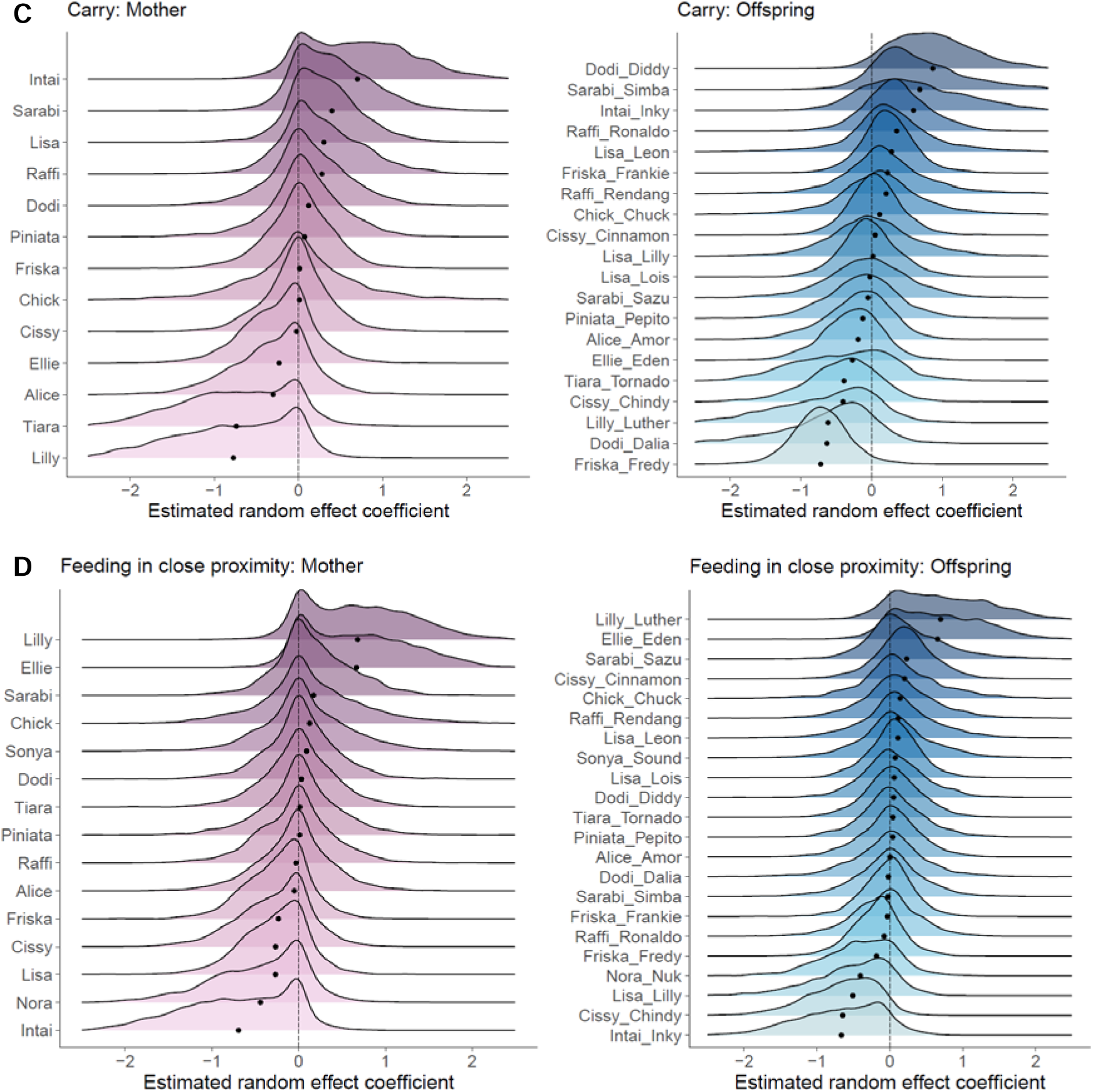
Between-individual variation in the average expression of four maternal behaviours after controlling for the confounding effects. Posterior distributions (on the logit scale) are shown for (A) *body contact termination* (random intercept model), (B) *close proximity termination* (full model), (C) *carrying* (full model), and (D) *feeding in close proximity* (full model) for the individual mothers and offspring relative to the estimated population-level average. Black dots show the posterior mean for each individual. Positive estimates indicate higher than average behavioural expression, while negative estimates indicate lower than average behavioural expression. Random intercepts of the mothers were calculated based on their behaviour with 1-3 offspring. Regardless of offspring of the same mother being similar to (e.g., *close proximity termination*: both Simba and Sazu being below population average) or different (e.g., *carrying*: Fredy being below and Frankie being above population average) from each other, mothers still showed consistent, significant variation in these behaviours.

**Table 1.**
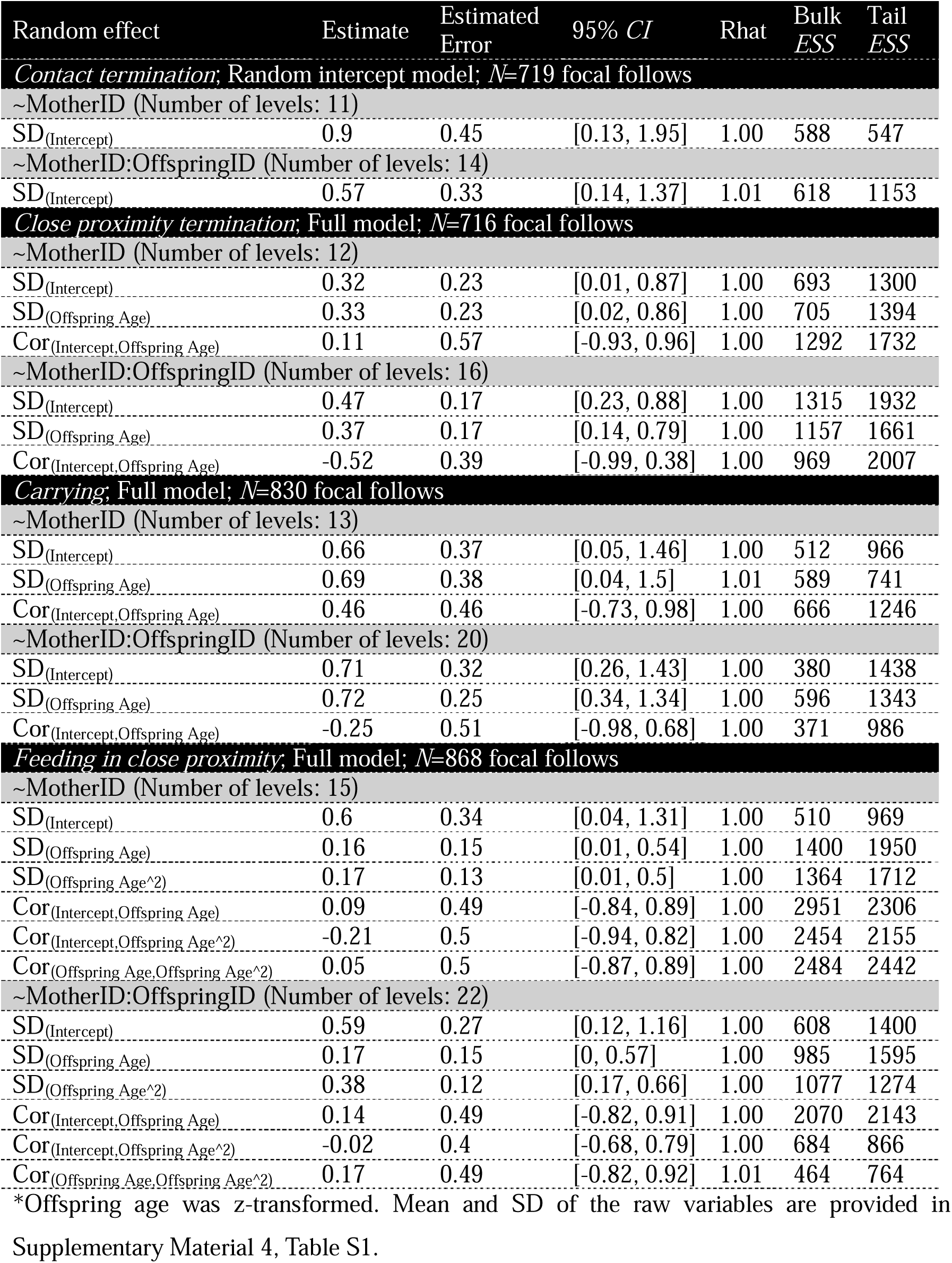
Estimated between-individual variation in the average expression of a behaviour and between-individual variation in behavioural plasticity. Means and standard deviations of the posterior distribution of parameters, their 95% credible intervals (*CI*), along with *ESS* for the mother and offspring random effects are shown for the four maternal behaviours in which there was significant between-individual variation among mothers. MotherID:OffspringID denotes nested random effect.

| Random effect | Estimate | Estimated Error | 95% CI | Rhat | Bulk ESS | Tail ESS |
| --- | --- | --- | --- | --- | --- | --- |
| **Contact termination; Random intercept model; N=719 focal follows** |  |  |  |  |  |  |
| ~MotherID (Number of levels: 11) |  |  |  |  |  |  |
| SD<sub>(Intercept)</sub> | 0.9 | 0.45 | [0.13, 1.95] | 1.00 | 588 | 547 |
| ~MotherID:OffspringID (Number of levels: 14) |  |  |  |  |  |  |
| SD<sub>(Intercept)</sub> | 0.57 | 0.33 | [0.14, 1.37] | 1.01 | 618 | 1153 |
| **Close proximity termination; Full model; N=716 focal follows** |  |  |  |  |  |  |
| ~MotherID (Number of levels: 12) |  |  |  |  |  |  |
| SD<sub>(Intercept)</sub> | 0.32 | 0.23 | [0.01, 0.87] | 1.00 | 693 | 1300 |
| SD<sub>(Offspring Age)</sub> | 0.33 | 0.23 | [0.02, 0.86] | 1.00 | 705 | 1394 |
| Cor<sub>(Intercept,Offspring Age)</sub> | 0.11 | 0.57 | [-0.93, 0.96] | 1.00 | 1292 | 1732 |
| ~MotherID:OffspringID (Number of levels: 16) |  |  |  |  |  |  |
| SD<sub>(Intercept)</sub> | 0.47 | 0.17 | [0.23, 0.88] | 1.00 | 1315 | 1932 |
| SD<sub>(Offspring Age)</sub> | 0.37 | 0.17 | [0.14, 0.79] | 1.00 | 1157 | 1661 |
| Cor<sub>(Intercept,Offspring Age)</sub> | -0.52 | 0.39 | [-0.99, 0.38] | 1.00 | 969 | 2007 |
| **Carrying; Full model; N=830 focal follows** |  |  |  |  |  |  |
| ~MotherID (Number of levels: 13) |  |  |  |  |  |  |
| SD<sub>(Intercept)</sub> | 0.66 | 0.37 | [0.05, 1.46] | 1.00 | 512 | 966 |
| SD<sub>(Offspring Age)</sub> | 0.69 | 0.38 | [0.04, 1.5] | 1.01 | 589 | 741 |
| Cor<sub>(Intercept,Offspring Age)</sub> | 0.46 | 0.46 | [-0.73, 0.98] | 1.00 | 666 | 1246 |
| ~MotherID:OffspringID (Number of levels: 20) |  |  |  |  |  |  |
| SD<sub>(Intercept)</sub> | 0.71 | 0.32 | [0.26, 1.43] | 1.00 | 380 | 1438 |
| SD<sub>(Offspring Age)</sub> | 0.72 | 0.25 | [0.34, 1.34] | 1.00 | 596 | 1343 |
| Cor<sub>(Intercept,Offspring Age)</sub> | -0.25 | 0.51 | [-0.98, 0.68] | 1.00 | 371 | 986 |
| **Feeding in close proximity; Full model; N=868 focal follows** |  |  |  |  |  |  |
| ~MotherID (Number of levels: 15) |  |  |  |  |  |  |
| SD<sub>(Intercept)</sub> | 0.6 | 0.34 | [0.04, 1.31] | 1.00 | 510 | 969 |
| SD<sub>(Offspring Age)</sub> | 0.16 | 0.15 | [0.01, 0.54] | 1.00 | 1400 | 1950 |
| SD<sub>(Offspring Age^2)</sub> | 0.17 | 0.13 | [0.01, 0.5] | 1.00 | 1364 | 1712 |
| Cor<sub>(Intercept,Offspring Age)</sub> | 0.09 | 0.49 | [-0.84, 0.89] | 1.00 | 2951 | 2306 |
| Cor<sub>(Intercept,Offspring Age^2)</sub> | -0.21 | 0.5 | [-0.94, 0.82] | 1.00 | 2454 | 2155 |
| Cor<sub>(Offspring Age,Offspring Age^2)</sub> | 0.05 | 0.5 | [-0.87, 0.89] | 1.00 | 2484 | 2442 |
| ~MotherID:OffspringID (Number of levels: 22) |  |  |  |  |  |  |
| SD<sub>(Intercept)</sub> | 0.59 | 0.27 | [0.12, 1.16] | 1.00 | 608 | 1400 |
| SD<sub>(Offspring Age)</sub> | 0.17 | 0.15 | [0, 0.57] | 1.00 | 985 | 1595 |
| SD<sub>(Offspring Age^2)</sub> | 0.38 | 0.12 | [0.17, 0.66] | 1.00 | 1077 | 1274 |
| Cor<sub>(Intercept,Offspring Age)</sub> | 0.14 | 0.49 | [-0.82, 0.91] | 1.00 | 2070 | 2143 |
| Cor<sub>(Intercept,Offspring Age^2)</sub> | -0.02 | 0.4 | [-0.68, 0.79] | 1.00 | 684 | 866 |
| Cor<sub>(Offspring Age,Offspring Age^2)</sub> | 0.17 | 0.49 | [-0.82, 0.92] | 1.01 | 464 | 764 |

\*Offspring age was z-transformed. Mean and SD of the raw variables are provided in Supplementary Material 4, Table S1.

For two other behaviours, *contact initiation* and *close proximity initiation*, the models’ predictive performance did not improve with the addition of random effects of mother identity (Supplementary Material 5, Table S1).

We found the largest estimated between-individual variation among mothers for the behaviours *contact termination* and *carrying* (Table 1, Figure 1A,C), while *close proximity termination* and *feeding in close proximity* had lower estimated between-individual variation among mothers (Table 1, Figure 1B,D). Furthermore, the estimated average expression of a behaviour for each mother varied across the behaviours (Figure 1A-D).

### Influence of confounding effects

When we compared the estimated between-mother variation in the average expression of maternal behaviour obtained from models in which we did and did not control for the known confounding effects, we found that for three of the analysed maternal behaviours not controlling for the confounding effects resulted in a higher estimated between-mother variation than controlling for these effects (Figure 2). In contrast, for three other behaviours, not controlling for the confounding effects reduced the estimated between-individual variation. The difference was most pronounced for *body contact initiation* and *termination*, where the models that did not control for the known confounding effects (Supplementary Material 3, Table S1) estimated the between-individual variation to be more than twice as high as the models that did control for the confounding effects (Figure 2). The differences between the estimated between-individual variations were moderate for *close proximity termination* and *feeding in close proximity* and negligible for *close proximity initiation* and *carrying*. Additionally, for three of the behaviours (i.e., *body contact initiation* and *termination*, and *close proximity termination*), the 95% credible intervals associated with the estimates were wider when the confounding effects were not controlled than when they were controlled for in the analysis (Figure 2). On the contrary, for the rest of the behaviours (i.e., *carrying*, *close proximity initiation*, and *feeding in close proximity*), the 95% credible intervals associated with the estimates were wider when the confounding effects were controlled than when they were not controlled for in the analysis (Figure 2).

**Figure 2.**
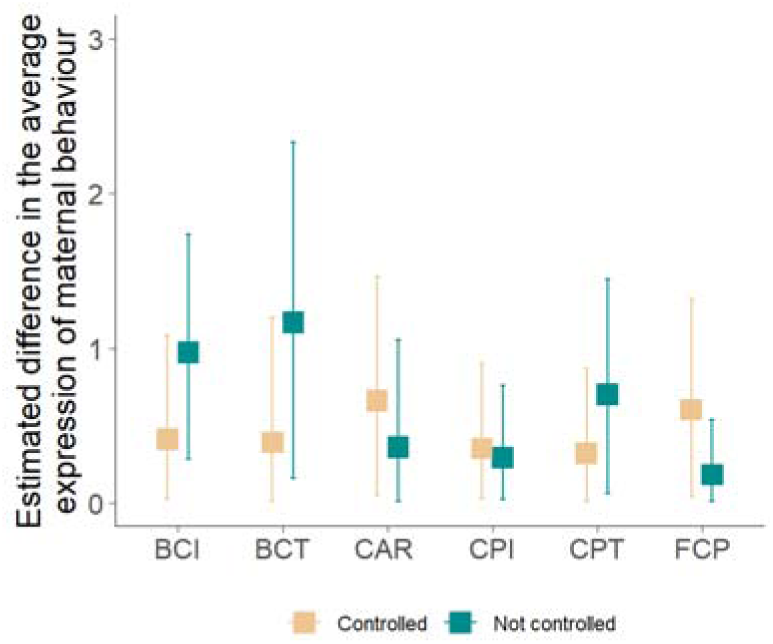
Comparison of the estimated differences among mothers in maternal behaviour. Estimated between-individual variation in the average expression of maternal behaviour (i.e., the estimated standard deviation of mother identity intercepts) is shown for each of the six maternal behaviours when the respective confounding effects were (full model) and were not controlled for in the analysis. The vertical lines represent 95% CI associated with the estimates. BCI: *Body contact initiation*; BCT: *Body contact termination*; CAR: *Carrying*; CPI: *Close proximity initiation*; CPT: *Close proximity termination*; FCP: *Feeding in close proximity*.

### Between-individual variation in maternal behavioural plasticity

For three of the behaviours – *close proximity termination, carrying*, and *feeding in close proximity* there was evidence of significant and consistent (i.e., across their different offspring) between-individual variation among mothers (SD: 0.16-0.69; Table 1) in behavioural plasticity over offspring age (Supplementary Material 5, Table S1) for these behaviours. Furthermore, offspring identity contributed to, but did not fully explain, the variation in maternal behavioural plasticity over offspring age (SD: 0.17-0.72). The largest between-individual variation in behavioural plasticity among mothers was observed for c*arrying* (Table 1, Supplementary Material 8, Figure S1). The significant effect of random slope of offspring age within mother identity suggests that mothers varied in how they adjusted their behaviour as their offspring developed, leading to mother-specific trajectories of maternal behaviour over offspring age (Figure 3).

**Figure 3.**
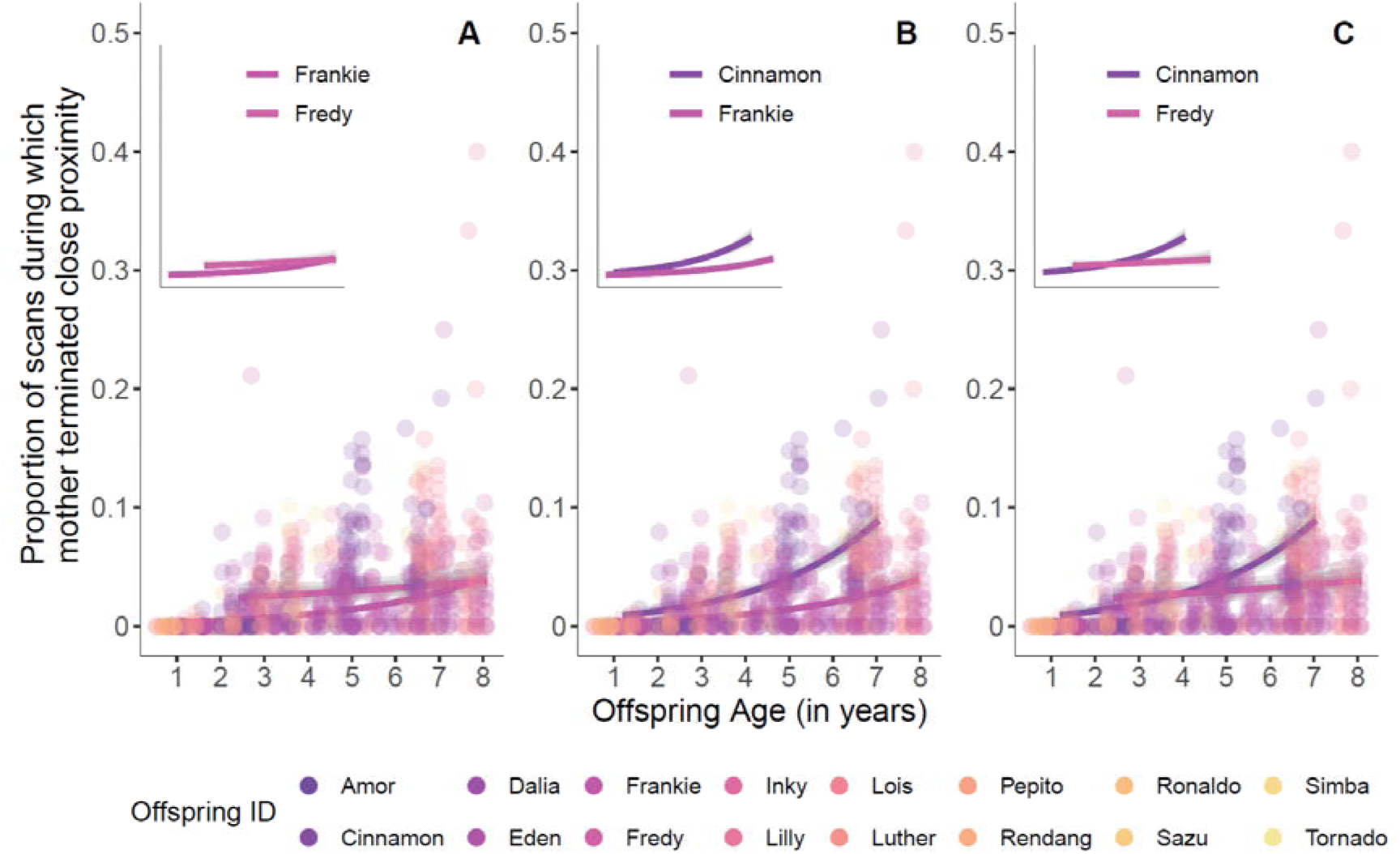
Consistent between-mother variation in behavioural plasticity. Fitted lines and their credible intervals (95%, 80%, and 50% in grey) are shown, as an example, for *close proximity termination* for A) two offspring of the same mother and B) & C) two offspring of two unrelated mothers from the full model to visualize the distinct trajectories of maternal behaviour over offspring age. Plots without the datapoints of all the focal individuals are shown in the insets for clearer visualization of the trajectories.

When we repeated the analysis with only those mothers with more than one offspring for *carrying* and *feeding in close proximity*, we again found significant and consistent between-mother variation in the average expression of as well as plasticity in maternal behaviour (Supplementary Material 9, Table S1 & S2, Figure S1 & S2).

## 4. Discussion

Much research to date on primate maternal behaviour has focused on the effect of biological characteristics of the mother and offspring, social factors, and ecological conditions on maternal behaviour [40,79–81], but little attention has been paid to individuals as a source of variation. Here we used a behavioural reaction norm approach to investigate between-individual variation in the average expression of maternal behaviour and between-individual variation in behavioural plasticity during offspring development in Sumatran orangutans. We found that mothers substantially differed in their average expression of maternal behaviour even after controlling for known confounding effects. Furthermore, we found that not controlling for the confounding effects either exaggerated or masked estimated between-individual variation. We further found that mothers differed in how they modified their behaviour in response to their offspring’s increasing age – i.e., mothers showed individualistic trajectories of maternal behaviour during offspring development and, thus, between-individual variation in behavioural plasticity. Importantly, we found that the identity of the offspring also contributed to variation among mothers in the average expression of maternal behaviour and plasticity in maternal behaviour over offspring age. However, maternal variation remained significant even after accounting for the offspring’s contribution, suggesting that mothers behave in a consistent manner across one’s different offspring [39].

As predicted, our analyses revealed substantial between-individual variation in the average expression of *carrying*, *feeding in close proximity*, *body contact termination*, and *close proximity termination* among mothers when controlling for confounding effects. This suggests that Sumatran orangutan mothers differ in the expression of these maternal behaviours regardless of the experienced socioecological conditions. Differences among mothers in maternal behaviours such as carrying, rejection, restraint, and ventral contact have been previously found in Guinea baboons, rhesus macaques, and vervet monkeys [13,26,39] – though these studies did not control for all likely confounding effects, leading to uncertainty in the extent of estimated variation. Furthermore, a recent study found individual differences in offspring-directed maternal communication in orangutans. However, this study included only one offspring per mother, making it impossible to tease apart the contribution of the mother *vs* the offspring to maternal behavioural variation [41]. Our results indicate that the differences in maternal behaviour are likely consistent in the mothers across their different offspring, strengthening the notion that there is personality – specifically maternal personality – in nonhuman primates [15]. In species with large brains, developmental effects that lead to consistent individual differences are likely to occur [82]. This is because the larger the brain (relative to body size) the more important developmental effects become, as large-brained species are more prone to developmental inputs [83], and a large share of the brain development happens after birth [84]. In Sumatran orangutans (also in our data set), *body contact initiation* and *close proximity initiation* with offspring are both rarely shown by mothers [40]. Accordingly, most of the data points for these behaviours in our dataset were zero, which makes it difficult to discern variation with the currently available analysis techniques.

In our study, the magnitude of variation among mothers was higher for *body contact termination* and *carrying* than that for *close proximity termination* and *feeding in close proximity*. In orangutans, *carrying* is a form of locomotory support [40,62] and close range feeding can provide social learning opportunities for offspring [61], whereas body contact and proximity maintenance are often associated with protection/thermoregulation in primates [85]. *Carrying* is energy intensive [86], and close range feeding may increase feeding competition and thus also have negative energetic effects for the mothers. Individual differences in the expression of a maternal behaviour could be the consequence of differences in the maximum investment that individual mothers can show (based on her genetic foundation or irreversible effects of early developmental experience). A mother’s genetically or developmentally determined body condition may thus constrain her investment in energetically critical maternal behaviour. However, since mother-offspring interactions are always dyadic in nature, it is possible that the differences among mothers may be at least partly driven by differences among their offspring. As predicted, we found non-zero variation among offspring of the same mother, suggesting that differences among offspring in their characteristics and demands can lead to differential expression of maternal behaviour. However, offspring explained only part of the variation in maternal behaviour.

Mothers differed in their average expression across the different behaviours. In primates, variation in maternal behaviours usually falls along two dimensions, namely protection and rejection [32,79], meaning that a mother’s behaviour in one dimension does not predict her behaviour in another dimension. Our results showed that orangutan mothers exhibit distinct individual patterns of investment across the different maternal behaviours. This dissimilarity suggests that the maternal behaviours that we analysed may fall on different dimensions in Sumatran orangutans, as that seen in captive chimpanzees and bonobos and many wild primates [22,32,79]. However, mothers who showed higher than average *carrying* showed significantly less than average *feeding in close proximity* with their offspring. In Sumatran orangutans, carrying is at its peak for the first two years of life and steeply declines thereafter, and feeding in proximity peaks between 5-6 years of age and then declines [40]. This suggests that early life investment in offspring may lead to faster feeding skill acquisition, resulting in a decrease in feeding in close proximity between the mother and the offspring later in life.

As predicted, we found that failing to appropriately control for the known confounding effects while estimating between-individual variation in the average expression of maternal behaviours can result in exaggerated estimation of variation among individuals. When differences in the biological characteristics and the socioecological environment under which the different mother-offspring pairs were observed [40] cause differences among individual behaviour, then by not deliberately adding these factors as predictors in their models, researchers risk wrongly assigning these differences to individual effects (i.e., creating false-positive between-individual differences) [26,43]. In addition, as predicted, our results also show that one may also end up underestimating individual variation (i.e., creating false-negative similarity between individuals) when not controlling for confounding effects. Failure to control for the confounding effects not only affects the estimated variation between mothers but also the uncertainty associated with it. Credible intervals were wider when confounding effects were not controlled for than when they were controlled for in half of our models. As the values of the different predictors, such as offspring age, association size, etc., change over time and as females are responding to the changes in these variables [40], differences in the effects of these variables further add on to the inherent differences among individuals, resulting in increased uncertainty around the estimates. In general, this result highlights the importance of controlling for the confounding effects in animal personality studies.

Though variation in behavioural plasticity is usually studied along an environmental or a social gradient [41,42,45,51], we investigated whether mothers differ in how they adjust their behaviour over offspring development. As predicted, first, our analyses revealed that offspring identity contributed to differences in behavioural plasticity among mothers. Second, our analyses revealed substantial variation in behavioural plasticity among mothers in response to offspring’s age in *close proximity termination, carrying*, and *feeding in close proximity*, even after accounting for the offspring’s contribution. This means that mothers differed in how they adjusted their behaviour during offspring development. This may be a result of some mothers being more plastic than others. As with between-individual difference in the average expression of these behaviours, because *body contact* and *close proximity initiation* are rarely shown by mothers, it is difficult to discern between-individual variation in plasticity in these two behaviours. Our analyses did not detect substantial between-individual variation in plasticity in *contact termination*, suggesting that the range of behavioural expression for *body contact termination* is indeed similar across mothers, despite the significant between-individual variation in the average expression.

Differences in mothers’ physiological conditions can lead to differences in their ability or willingness to respond to their offspring’s needs, which may be a source of individual differences in plasticity in maternal behaviour. In line with this reasoning, we found the most pronounced between-individual variation in plasticity for the most energetically costly behaviour, namely *carrying*. Maternal condition was also found to influence between-individual differences in time spent in and out of contact with offspring and frequency of rejection in other primates [81]. However, as evidence suggests that orangutan mothers are capable of adjusting their behaviour in response to their offspring’s age [40,60], it is possible that variation in the development of the offspring’s locomotor skill levels lead to differences in the mothers’ readiness to carry them. In other words, differences in offspring’s pace of development and, especially, variation in the magnitude of these differences can bring about individual differences in maternal behavioural plasticity. Additionally, differences among offspring in their health could drive differences in maternal behaviour, however, all focal offspring in this study were in apparent good health, and none of them died during or after the study (consistent with published extremely low infant mortality; [57]).

Overall, our results support that there is consistent between-individual behavioural variation in maternal behaviour in Sumatran orangutans. However, as with any field-based study, individual mothers differed in the conditions under which they were observed, which may affect both individual differences in the average expression and plasticity of the maternal behaviours. Even though we controlled for previously established confounding effects [40], there may be other biological state characteristics or socioecological variables that affect variation in maternal behaviour that we currently cannot quantify or are not even aware of. Furthermore, the complexity of these effects may exceed the way we quantified them. For example, we took into account the current presence or absence of males because of their known effects on maternal behaviours, which likely arise from the risk they may pose to immature individuals [40,87]. However, individual males likely differ in the risk they pose to the offspring [88] and thus also in the effects they have on maternal behaviour. Controlling for more accurately quantified prevailing social risk faced by offspring would thus be a better approach, even though not feasible with the current dataset. Despite our study making use of around 6000 hours of behavioural observation conducted over the span of 15 years, it is constrained by the limited number of focal individuals, the offspring’s lengthy developmental periods, and long inter-birth intervals. Accordingly, sample size and sampling scheme substantially varied among the mothers, which inevitably limits the inferences possible with our study. For a more accurate quantification of between-individual differences in maternal behaviour, we need data on mother-offspring pairs spanning the entire range of offspring developmental periods and, ideally, data on multiple offspring for all the mothers.

Our study aimed to disentangle actual pattern from noise through partitioning observed variation in maternal behaviour. Our results suggest that Sumatran orangutan maternal behaviour is shaped by individuality, apart from mother-offspring social and biological state characteristics and socioecological conditions [40]. Our study adds to previous research on maternal personality in primates and other species in showing that there are consistent differences among mothers in their behaviour towards offspring [13,26,32,39,48,66]. Our study signifies the importance of expanding research focus from average behaviour to individual behaviour such that variation around the mean is not considered as pure noise. Consistent between-individual variation in the average expression of a behaviour and plasticity can have important consequences and present novel opportunities for studies of the evolutionary processes that shape the behaviour.

For consistent, heritable variation to undergo natural selection, there should also be fitness consequences associated with such variation. Investigating whether differences in maternal behaviour are associated with differences in the speed at which offspring reach developmental milestones and the offspring’s future reproductive success will shed light on the adaptive value of maternal behaviour for both the mother and the offspring. Our study also sets the stage to address whether daughters inherit maternal personality – socially and/or biologically – potentially giving rise to matrilineal differences in maternal behaviour [13,63]. Ultimately, studies on individual variation in maternal behaviour can help shed light on the causes, maintenance, and consequences of such variation.

## Ethics

As orangutans are arboreal and as observers are on the ground, the minimum observation distance was at least 7 m in the field. This minimises the effect of observer presence on the natural behaviour of the individuals. Data collection was purely observational in nature, and non-invasive. The research followed the recommendations of the Animals (Scientific Procedures) Act 1986, as published by the UK government, and the principles of Ethical Treatment of Non-Human Primates as stated by the American Society of Primatologists. Approval for this study was obtained from the Indonesian State Ministry for Research and Technology (RISTEK).

## Data accessibility

Data and original code are deposited at Dryad data repository.

## Authors’ contributions

Conceptualization: TR, MvN, MF, CS; Data collection: TUN, CS; Data curation: TR, CS; Methodology: TR, RM, PB, CS; Formal analysis: TR; Funding acquisition: CS; Investigation: TR; Field project administration: SSUA, CS; Supervision: CS; Visualization: TR; Writing—original draft: TR; Writing—review & editing: TR, RM, SSUA, TUN, MvN, MF, PB, CS.

## Acknowledgements

We thank all the dedicated field staff, students, and research assistants – in particular, Armas Fitra, Adami Khairul, Fikar Zulkarnean, Julia Kunz, Lara Nellissen, Mudin, Natasha Bartolotta, Saidi Agan, Toni M. Sunil, and Ulil Azhari – who, besides Tazkia Umaira Aprilla and Caroline Schuppli, collected most of the data used in this study. We are grateful to the Indonesian State Ministry for Research and Technology (BRIN-RISTEK), the Directorate General of Natural Resources and Ecosystem Conservation–Ministry of Environment and Forestry of Indonesia (KSDAEKLHK), the Ministry of Internal Affairs, Indonesia, the Sumatran orangutan conservation Program (SOCP), Balai Besar Taman Nasional Gunung Leuser (TNGL), in particular Arif Saifudin, Zakir, and Samsul Amar, and all staffs in Medan for their permission and support to conduct this study. We thank Universitas Nasional (UNAS) for their support and collaboration. We thank Tri Rahmaeti for coordinating our research permits and liaising with Indonesian offices and institutions. We thank the technical assistants, Richard Young and Francois Lamarque, for maintaining the SUAQ database. Sumatran orangutan mothers differ in the extent and trajectory of their expression of maternal behaviour

