## Supplemental Material for "Sumatran orangutan mothers differ in the extent and trajectory of their expression of maternal behaviour"

**Supplementary Material 1**

*Behavioural Reaction Norms*

**Box 1: Behavioural Reaction Norms**

With the behavioural reaction norm approach [40,42–45], repeated observations of individuals over time are analysed to divide the observed behavioural variance into fixed effects and random effects (including within- and between- individual variation) components using a regression method. The intercept of the model represents the population’s average expression of a behaviour. Behavioural plasticity is inferred via the slope of the regression, which represents behavioural variation over a predictor’s range at the population level. Between-individual variation in the average expression of a behaviour is measured as the difference between the population-level and the individual-level intercepts. Between-individual variation in behavioural plasticity is measured as the difference between the population-level and individual-level slopes.

*Maternal behaviour*

**Supplementary Material 1, Table S1**. Maternal behaviours that were analysed in the study, and the numerator and denominator used to calculate maternal behaviours based on [1]. For each behaviour, the opportunities available to the mothers to show that behaviour were controlled for in the denominator.

| **Behaviour** | **Numerator** | **Denominator** |
| --- | --- | --- |
| Body contact initiation | No. of scans in which the mother initiated body contact with her offspring (>0 m to 0 m) | Total no. of scans in which the mother and her offspring were not in contact (i.e., >0 m) during a follow |
| Body contact termination | No. of scans in which the mother terminated body contact with her (0 m to >0 m) | Total no. of scans in which the mother and offspring were in contact during a follow |
| Close proximity initiation | No. of scans in which the mother initiated close proximity with her offspring (>5-<=50 m to <=5 m but no body contact) | Total no. of scans in which the mother and offspring were not in close proximity (i.e., >5-<=50 m) during a follow |
| Close proximity termination | No. of scans in which the mother terminated close proximity with her offspring (<=5 m, but no contact to >5-<=50 m) | Total no. of scans in which the mother and offspring were in close proximity (i.e., <=5 m, but no contact) during a follow |
| Carry | No. of scans in which the mother carried her offspring | Total no. of scans in which the mother moved during a follow |
| Feeding in close proximity | No. of scans in which the mother was feeding within 5 m of her offspring who was also feeding on the same food item as its mother | Total no. of scans during which the mother was feeding during a follow |

Though nursing is an important maternal behaviour, it often occurs inconspicuously in orangutans due to their arboreal lifestyle. Thus, our sample size did not allow to us to analyse this behaviour in our study. Maternal behaviours common to most primates such as grooming, social play, food sharing, and agonistic protection were only rarely observed in the study population (field data).

The arm span of an adult orangutan is around 2.2 m. Most feeding trees at Suaq have a crown diameter of more than 5 m, meaning that when in a feeding patch both mother and offspring can control the distance to each other. Mothers control the distance to their offspring in feeding trees by chasing their offspring away if they come too close, and they retrieve their offspring if they wander off too far [2].

**Supplementary Material 2**

*Sample size*

**Supplementary Material 2, Table S1**. Overview of the focal mothers and their offspring included in the current study.

| SI. No. | Mother  ID | Offspring  ID | Offspring  sex | Offspring DOB | Range of no. of follows of the mother-offspring pair (across the six behaviours) | No. of  behaviours sampled |
| --- | --- | --- | --- | --- | --- | --- |
| 1 | Alice | Amor | Male | 01-Jan-15 | 8-17 | 6 |
| 2 | Cissy | Chindy | Female | 01-Jan-03 | 12-13 | 2 |
| 3 | Chick | Chuck | Male | 01-Jul-07 | 5-5 | 2 |
| 4 | Cissy | Cinnamon | Female | 01-Apr-12 | 81-112 | 6 |
| 5 | Dodi | Dalia | Female | 01-Oct-13 | 5-6 | 4 |
| 6 | Dodi | Diddy | Male | 01-Dec-05 | 7-7 | 2 |
| 7 | Ellie | Eden | Female | 01-Nov-14 | 141-166 | 6 |
| 8 | Friska | Frankie | Male | 01-Aug-12 | 114-165 | 6 |
| 9 | Friska | Fredy | Male | 01-Jun-05 | 33-60 | 6 |
| 10 | Intai | Inky | Male | 01-Jul-03 | 6-8 | 6 |
| 11 | Lisa | Leon | Male | 15-Nov-18 | 19-21 | 2 |
| 12 | Lisa | Lilly | Female | 01-Mar-01 | 12-26 | 6 |
| 13 | Lisa | Lois | Male | 01-Aug-10 | 135-144 | 6 |
| 14 | Lilly | Luther | Male | 01-Mar-16 | 17-26 | 6 |
| 15 | Nora | Nuk | Male | 01-Jan-06 | 8-8 | 1 |
| 16 | Piniata | Pepito | Male | 01-Jan-13 | 10-12 | 6 |
| 17 | Raffi | Rendang | Male | 15-Jul-13 | 16-17 | 5 |
| 18 | Raffi | Ronaldo | Male | 01-Jan-06 | 8-13 | 5 |
| 19 | Sarabi | Sazu | Male | 01-Jul-06 | 15-17 | 4 |
| 20 | Sarabi | Simba | Male | 01-Mar-13 | 10-18 | 5 |
| 21 | Sonya | Sound | Female | 01-Aug-18 | 6-6 | 1 |
| 22 | Tiara | Tornado | Male | 01-Jul-14 | 15-19 | 6 |

| (a)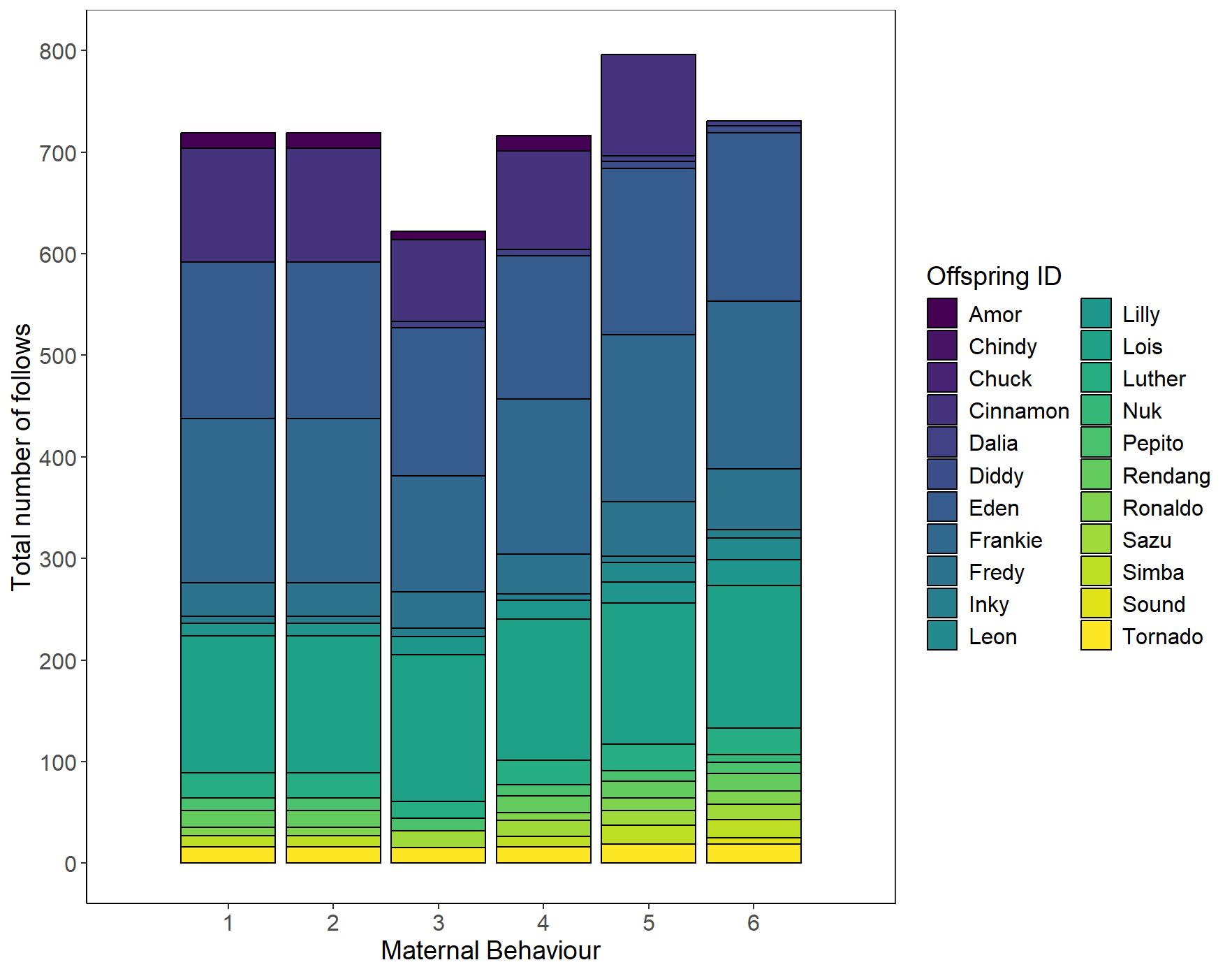 | (b)  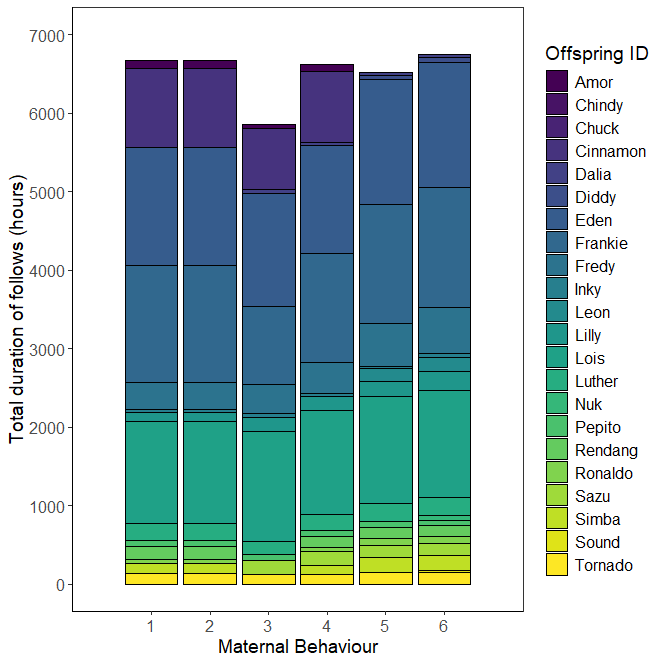 |
| --- | --- |

**Supplementary Material 2, Figure S1**. Total (a) number of follows and (b) duration of follows for each of the focal offspring for the six maternal behaviours analysed in the current study. The maternal behaviours are 1. body contact initiation, 2. body contact termination, 3. close proximity initiation, 4. close proximity termination, 5. carrying, and 6. feeding in close proximity.

To get reliable results, we used a cut-off of at least five follows per offspring per behaviour. Therefore, not all the offspring could be included in all the behavioural analyses. Despite these constraints, we had data on most behaviours for most offspring (Supplementary Material 2, Table S1). The number of mothers for whom we had sufficient data (i.e., at least 5 focal follows per offspring) on multiple offspring varied across the behaviours (*body contact initiation* and *termination*: 3/11 mothers; *close proximity initiation*: 2/11 mothers; *close proximity termination*: 4/12 mothers*; carrying*: 6/13 mothers; *feeding in close proximity*: 6/15 mothers).

*Sampling scheme*

| (a) Contact initiation  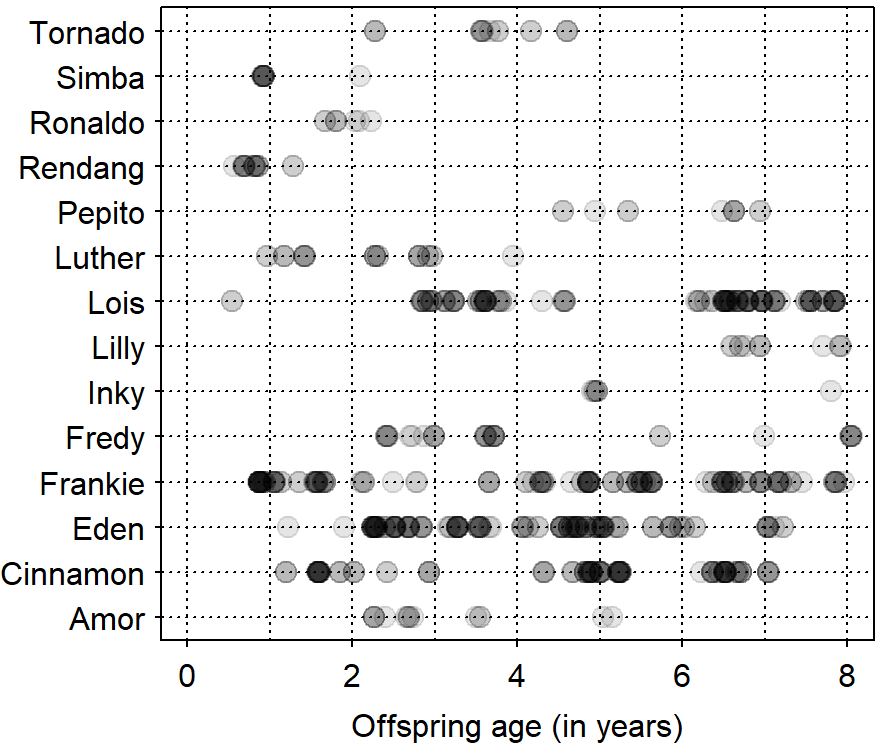 | (b) Contact termination  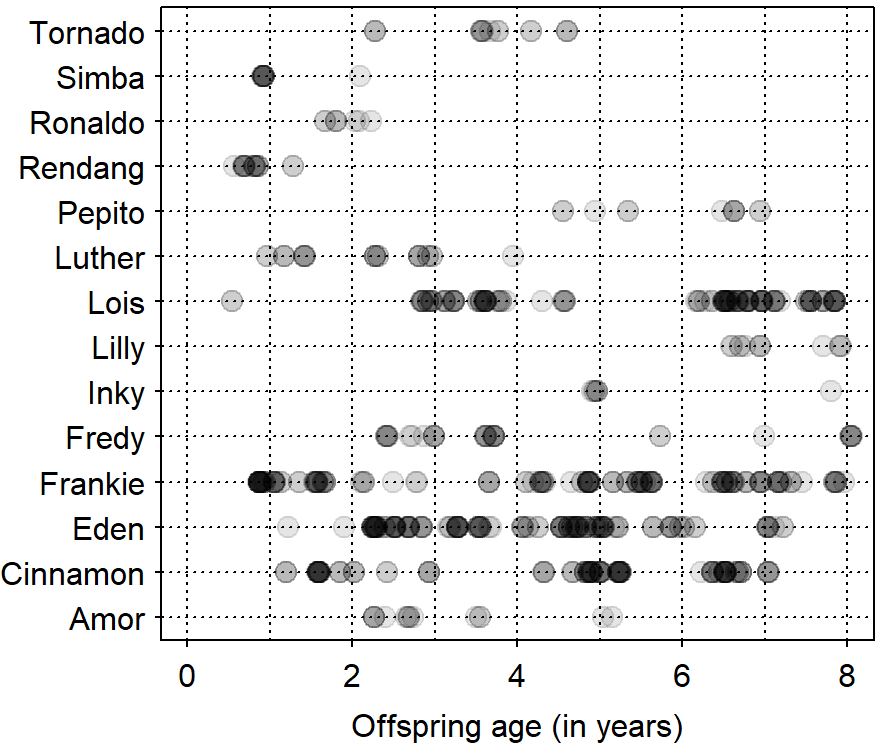 |
| --- | --- |
| (c) Close proximity initiation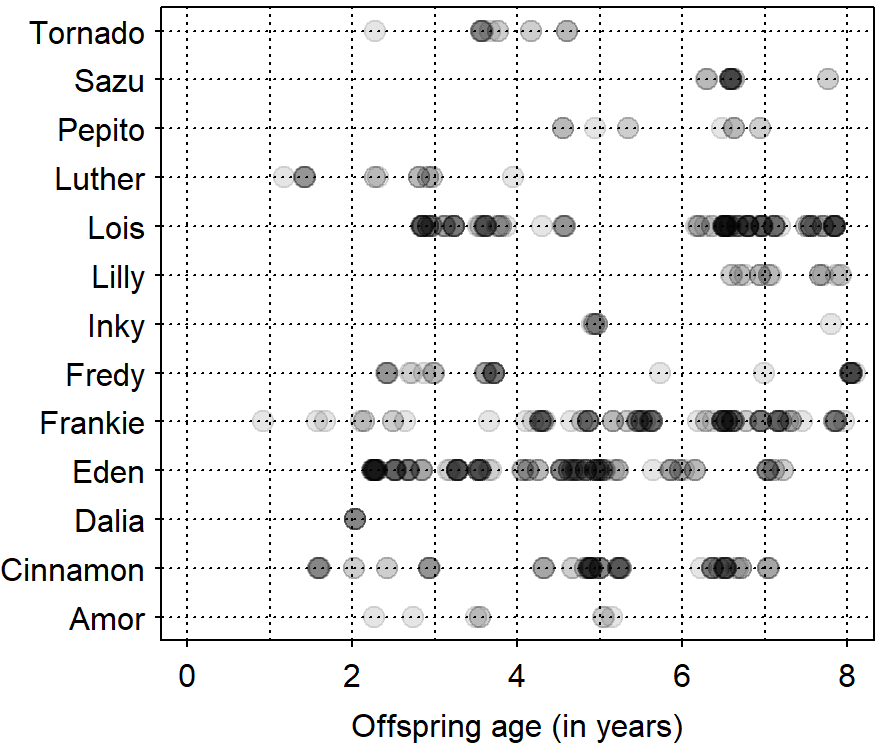 | (d) Close proximity termination  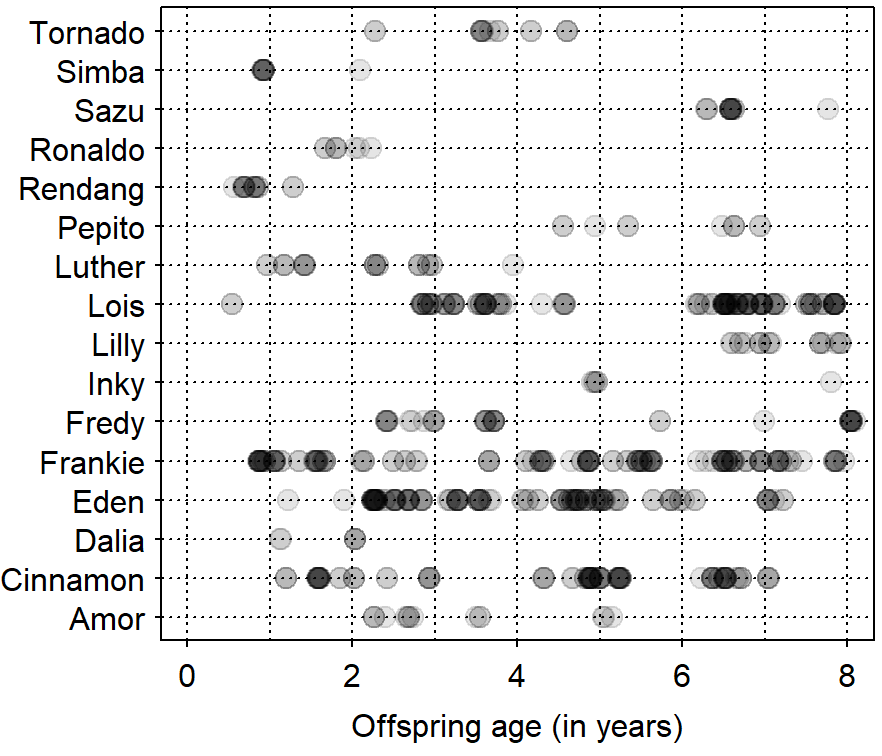 |
| (f) Carrying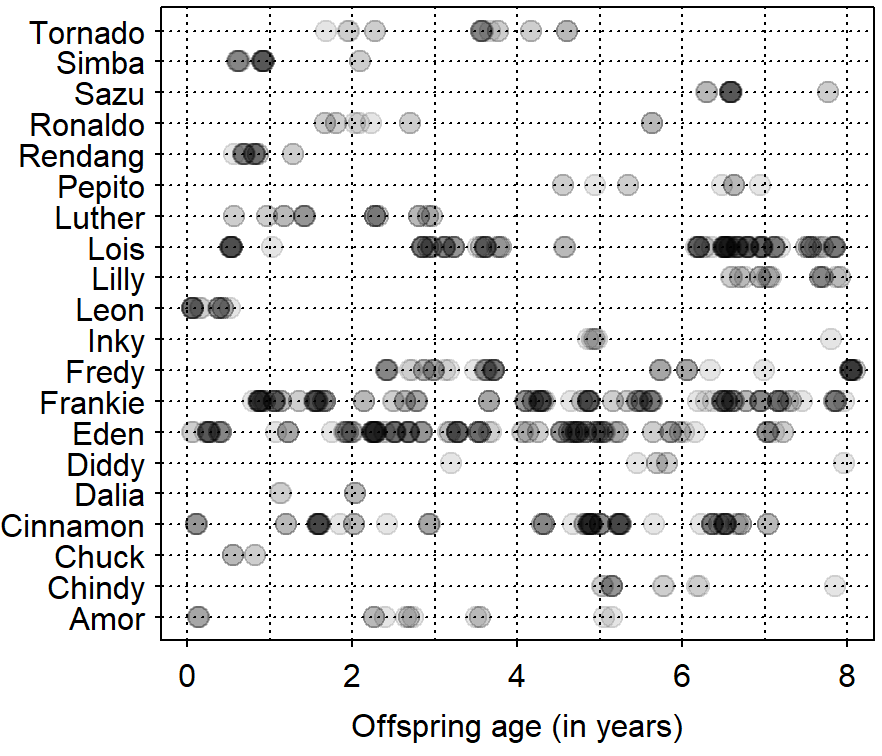 | (g) Feeding in close proximity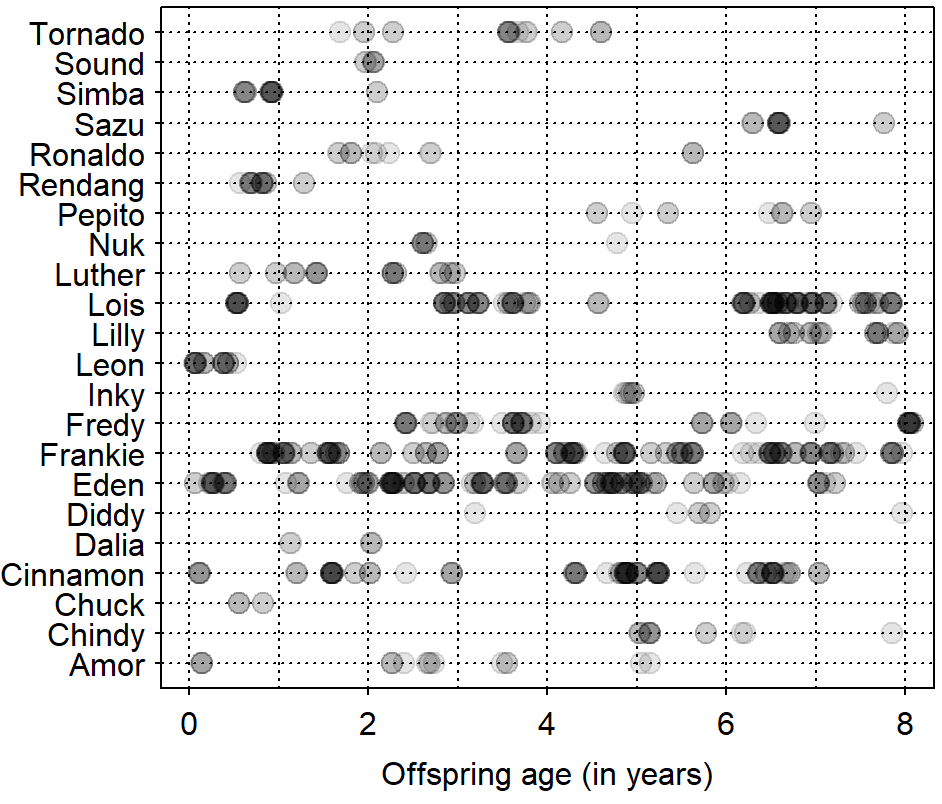 |

**Supplementary Material 2, Figure S2**. Overview of the sampling scheme, after the application of data filter for number of follows per offspring (see Methods for details), for each of the six behaviours of interest. Each filled circle in grey represents a focal follow, between birth and 8 years of age, during which the expression of a specific maternal behaviour could be assessed (note that we calculated the expression of each behaviour while controlling for the opportunities for it to be expressed, see Table S1). The filled dots appear darker proportionate to the number of focals recorded during a specific offspring age, as the dots overlap.

**Supplementary Material 3**

*Significant predictors of maternal behaviours in wild Sumatran orangutans*

**Supplementary Material 3, Table S1**. Effect of mother-offspring characteristics and socioecological factors on the six behaviours of interest from the study on the predictors of maternal behaviours in Sumatran orangutans [1]. NS indicates a predictor being not significant and * indicates a predictor being significant for a behaviour. (1) refers to mother-offspring characteristics and (2) socioecological factors.

| Maternal behaviour | Mother’s parity (1) | Offspring age (1) | Offspring sex (1) | FAI (2) | Average association size (2) | Male presence / absence (2) |
| --- | --- | --- | --- | --- | --- | --- |
| Body contact initiation | NS | * | NS | NS | * | NS |
| Body contact termination | NS | * | NS | NS | NS | NS |
| Close proximity initiation | NS | * | NS | NS | NS | NS |
| Close proximity termination | NS | * | NS | NS | * | NS |
| Carrying | NS | * | NS | NS | NS | * |
| Feeding in close proximity† | NS | * | * | * | NS | NS |

† Included both a linear and a quadratic effect of offspring age. Both the age effects were significant.

Mother’s parity was a categorical predictor with primiparous (females with their first offspring) and multiparous (females with more than one offspring) as the two levels. Offspring age was a continuous predictor and ranged between birth and eight years. Offspring sex was categorical with male and female as the two levels. Fruit availability index (FAI) was a continuous predictor and was calculated as the percentage of fruiting trees in the phenology transects per month in the study site. Average association size was a continuous predictor and was calculated as the number of individuals of any age-class within a 50 m distance of the focal individual, averaged across the two-minute scan samples during a focal follow. Male presence/absence was a categorial predictor with presence (a male in association with the focal individual) and absence (no male in association with the focal individual) as its two levels [see 1 for further details of all the predictors].

**Supplementary Material 4**

*Fixed effects or population-level effects*

**Supplementary Material 4, Table S1**. Results of the Bayesian regression obtained from the more parsimonious model (Supplementary Material 5, Table S1) for each of the six maternal behaviours. Mean values of the posterior distribution of parameter estimates, their 95% credible intervals (*CI*), along with Rhat and *ESS* for the fixed effects are shown.

| Fixed effect | Estimate | Estimated Error | 95% CI | Rhat | Bulk  ESS | Tail  ESS |
| --- | --- | --- | --- | --- | --- | --- |
| *Body contact initiation*; Fixed effects model; *N*=719 focal follows; *R*^2^=0.32 | | | | | |  |
| Intercept | -5.25 | 0.09 | [-5.44, -5.07] | 1.00 | 1003 | 1480 |
| Offspring Age | -1.11 | 0.08 | [-1.27, -0.97] | 1.00 | 1178 | 1512 |
| Association size | 0.27 | 0.05 | [0.17, 0.36] | 1.00 | 1706 | 1576 |
| *Body contact termination*; Random intercept model; *N*=719 focal follows; *R*^2^=0.48 | | | | | |  |
| Intercept | -2.37 | 0.36 | [-3.11, -1.69] | 1.00 | 1933 | 1965 |
| Offspring Age | 0.44 | 0.05 | [0.34, 0.54] | 1.00 | 5339 | 2222 |
| *Close proximity initiation*; Fixed effects model; *N*=622 focal follows; *R*^2^=0.31 | | | | | |  |
| Intercept | -3.65 | 0.05 | [-3.75, -3.54] | 1.00 | 1401 | 1578 |
| Offspring Age | -0.28 | 0.05 | [-0.39, -0.18] | 1.00 | 1205 | 1372 |
| *Close proximity termination*; Full model; *N*=716 focal follows; *R*^2^=0.40 | | | | |  |  |
| Intercept | -3.75 | 0.21 | [-4.17, -3.34] | 1.00 | 1546 | 1695 |
| Offspring Age | 0.71 | 0.21 | [0.35, 1.16] | 1.00 | 1914 | 1682 |
| Association size | -0.03 | 0.05 | [-0.12, 0.06] | 1.00 | 5138 | 2159 |
| *Carrying*; Full model; *N*=830 focal follows; *R*^2^=0.80 | | | |  |  |  |
| Intercept | -1.24 | 0.31 | [-1.87, -0.64] | 1.00 | 1714 | 1718 |
| Offspring Age | -2.13 | 0.32 | [-2.77, -1.52] | 1.00 | 1784 | 1846 |
| Male_P_A [P] | 0.07 | 0.08 | [-0.08, 0.23] | 1.00 | 3711 | 2093 |
| *Feeding in close proximity*; Full model; *N*=868 focal follows; *R*^2^=0.40 | | | | |  |  |
| Intercept | -1.73 | 0.42 | [-2.63, -0.91] | 1.00 | 2318 | 2113 |
| Offspring Age | 0.82 | 0.14 | [0.5, 1.06] | 1.01 | 742 | 1338 |
| Offspring Age^2 | -0.35 | 0.15 | [-0.65, -0.07] | 1.00 | 2410 | 2362 |
| Offspring Sex [Male] | 0.51 | 0.46 | [-0.38, 1.44] | 1.00 | 1821 | 2246 |
| FAI | -0.18 | 0.04 | [-0.26, -0.1] | 1.00 | 5490 | 2438 |

Mean ± SD of original (untransformed) variables for each behaviour:

Contact initiation: Offspring Age (4.340 ± 2.100), Association size (1.089 ± 1.081)

Contact termination: Offspring Age (4.340 ± 2.100)

Close proximity initiation: Offspring Age (5.064 ± 1.819)

Close proximity termination: Offspring Age (4.469 ± 2.137), Association size (0.840 ± 1.978)

Carrying: Offspring Age (4.079 ± 2.334)

Feeding in close proximity: Offspring Age (4.055 ± 2.322), FAI (10.127 ± 2.862)

We z-standardized the continuous predictors to a mean of zero and a standard deviation of one to ease model convergence and interpretation, and we dummy-coded and mean-centered offspring sex and male presence/absence, such that the intercept corresponds to the expected average maternal behavioural expression without being dependent on any one level of the categorical predictors. We used brms’ default (un-informative) priors. We ran one chain for each model with 5000 iterations, with a warm-up period of 2000 iterations. This resulted in an effective sample size (*ESS*) of >400 and an Rhat of 1 for each estimated parameter across all the models. To prevent divergent transitions, adapt-delta was set to 0.99.

| (a) Contact initiation  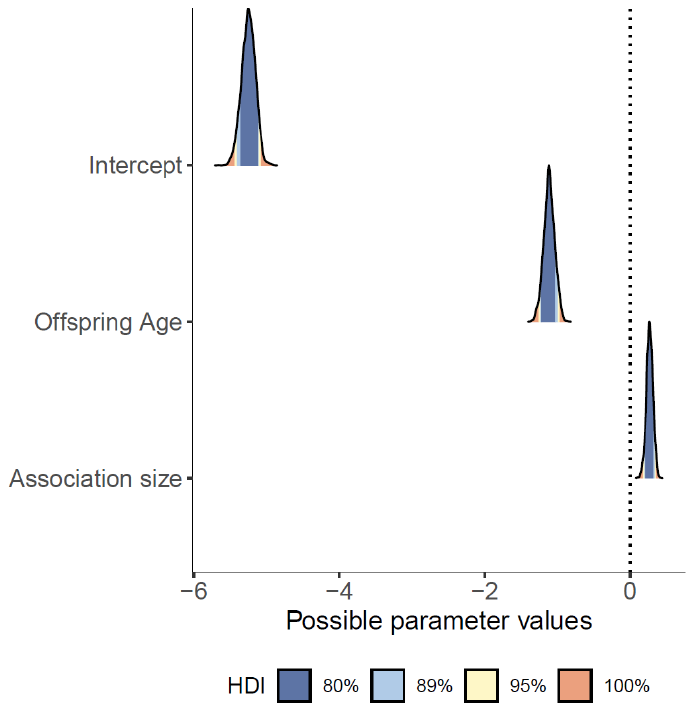 | (b) Contact termination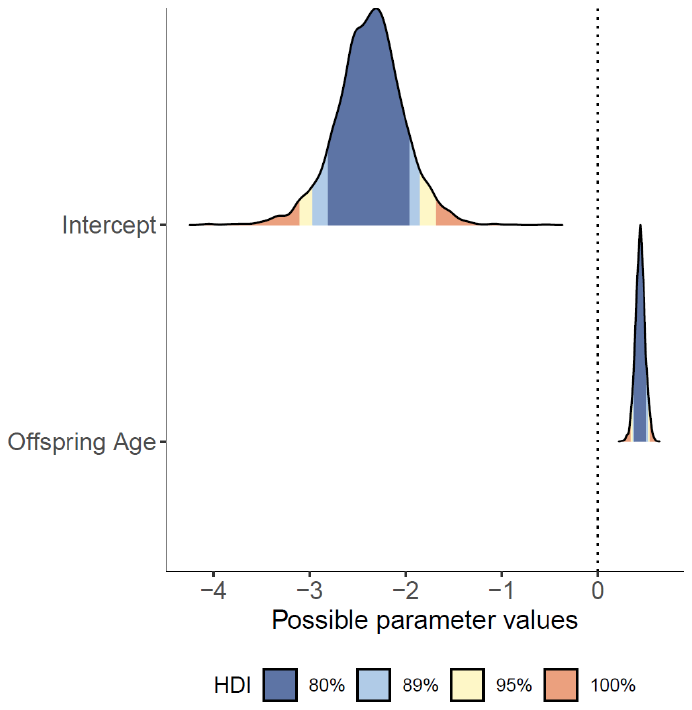 |
| --- | --- |
| (c) Close proximity initiation  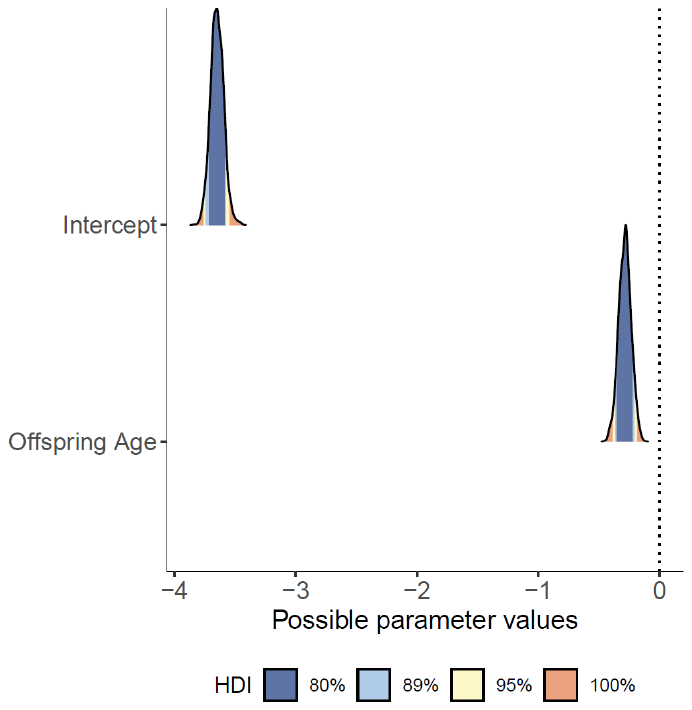 | (d) Close proximity termination  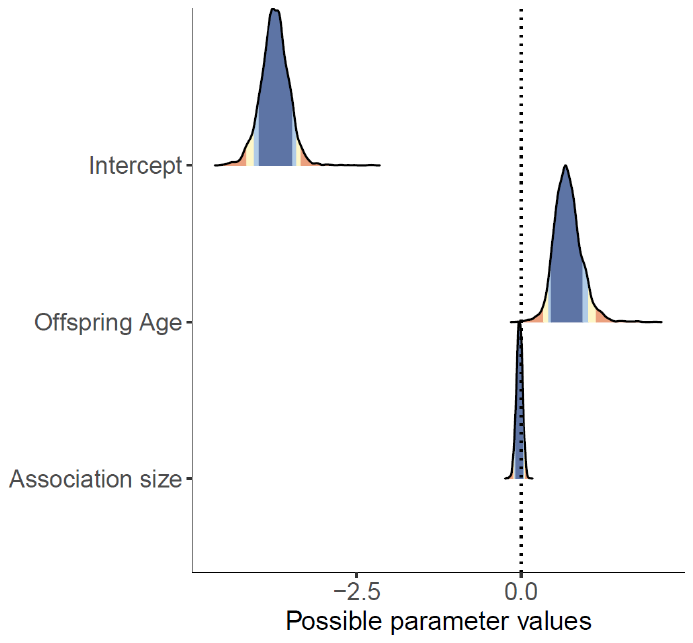 |
| (f) Carrying  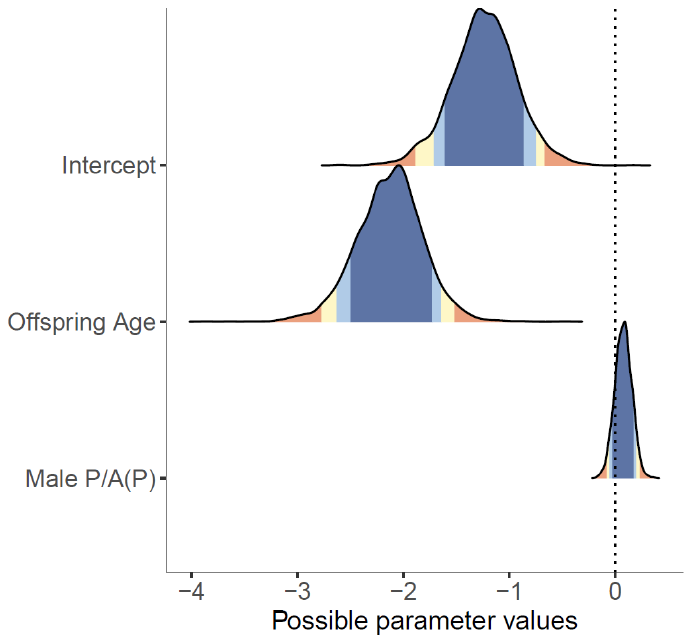 | (g) Feeding in proximity  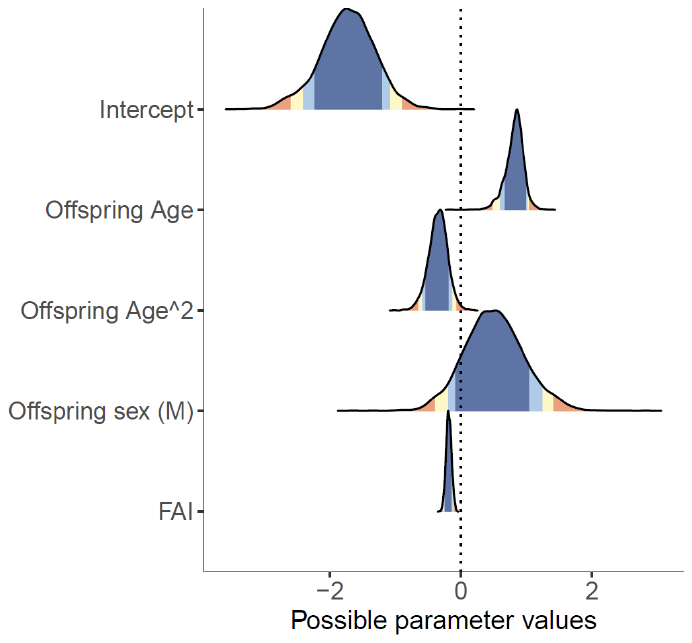 |

**Supplementary Material 4, Figure S1.** Highest Density Intervals (HDI) of the posterior distributions of the estimated population-level parameters from the more parsimonious model (Supplementary Material 4, Table S1) for each of the six maternal behaviours.

**Supplementary Material 5**

*LOO-CV model comparisons*

**Supplementary Material 5, Table S1**. Difference in predictive performance of the fitted models. Results of the model comparisons based on LOO-CV are shown. The most parsimonious model is marked with an asterisk (*). Δ*ELPD* ± 2 × Δ*SE* for each model was obtained by comparing it with the full model.

| Model | *ELPD_LOO* | *SE* | Δ*ELPD* | Δ*SE* | Δ*ELPD* ± 2 × Δ*SE* |
| --- | --- | --- | --- | --- | --- |
| ***Contact initiation*** | | | | | |
| Full model | -908.5 | 36.3 | 0 | 0 | - |
| Random intercept model | -914.3 | 36.2 | -5.8 | 3.8 | [-13.4, 1.8] |
| Fixed effects model* | -921.9 | 36.2 | **-13.4** | **6.9** | **[-27.2, 0.4]** |
| ***Contact termination*** | | | | | |
| Full model | -1443.4 | 29.9 | 0 | 0 | - |
| Random intercept model* | -1450.2 | 29.8 | **-6.9** | **5** | **[-16.9, 3.1]** |
| Fixed effects model | -1492.4 | 29.7 | -49.1 | 10 | [-69.1, -29.1] |
| ***Close proximity initiation*** | | | | | |
| Full model | -983.0 | 27.9 | 0 | 0 | - |
| Random intercept model | -983.1 | 28.1 | -0.1 | 1.9 | [-3.9, 3.7] |
| Fixed effects model* | -992.7 | 28.4 | **-9.7** | **5.3** | **[-20.3, 0.9]** |
| ***Close proximity termination*** | | | | | |
| Full model* | -1510.9 | 35.1 | **0** | **0** | - |
| Random intercept model | -1524.7 | 34.9 | -13.7 | 5.1 | [-23.9, -3.5] |
| Fixed effects model | -1565.8 | 35.9 | -54.9 | 10 | [-74.9, -34.9] |
| ***Carry*** | | | | | |
| Full model* | 2039.0 | 54.1 | **0** | **0** | - |
| Random intercept model | 1985.1 | 53.7 | -53.9 | 8.9 | [-71.7, -36.1] |
| Fixed effects model | 1939.7 | 54.0 | -99.2 | 12.3 | [-123.8, -74.6] |
| ***Feeding in close proximity*** | | | | | |
| Full model* | 1377.4 | 63.9 | **0** | **0** | **-** |
| Random intercept model | 1354.0 | 62.6 | -23.4 | 7.5 | [-38.4, -8.4] |
| Fixed effects model | 1255.2 | 62.5 | -122.2 | 16.6 | [-155.4, -89] |

Since our models were already too complex given the dataset and as slope estimates were zero for FAI, average association size, and male presence during a follow for most of the behaviours when analysed with a similar dataset [40], we did not include these variables in the random slopes part.

Leave-one-out cross-validation LOO-CV allowed us to examine whether the out-of-sample predictive performance of a model is improved by the addition of model terms (here, the random intercepts and slopes). It does so by estimating the difference in the expected log predictive density (*ELPD*). We considered two models to be similar in their predictive performance if Δ*ELPD* ± 2 × Δ*SE* overlapped zero, where Δ*ELPD* indicates the *ELPD* difference of both models and Δ*SE* the corresponding standard error.

If Δ*ELPD* ± 2 × *SE* between the full model and the fixed effects model overlapped zero, we inferred that adding the random effects of mother identity did not improve the predictive performance of the model (i.e., that our data does not provide convincing evidence that there was variation across mothers or across mothers over offspring age). Similarly, if Δ*ELPD* ± 2 × *SE* between the full model and the random intercept model overlapped zero, we inferred that adding the random slope of offspring age within the random intercept did not improve the predictive performance of the model (i.e., our data does not provide convincing evidence that there was variation across mothers over offspring age). We chose the more parsimonious model whose Δ*ELPD* ± 2 × *SE* did not overlap zero to draw inferences about between-individual differences in maternal behaviour.

**Supplementary Material 6**

*Posterior distribution of the estimated standard deviation for the random intercept of mother identity and offspring identity nested within mother identity*

| (a) *Contact termination*  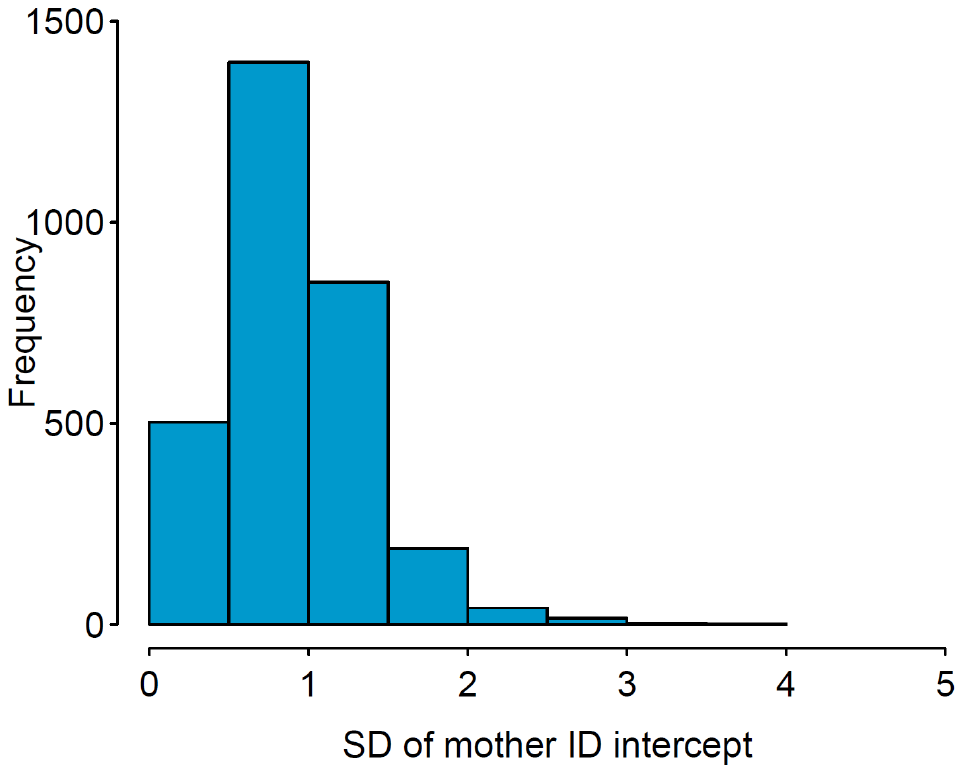  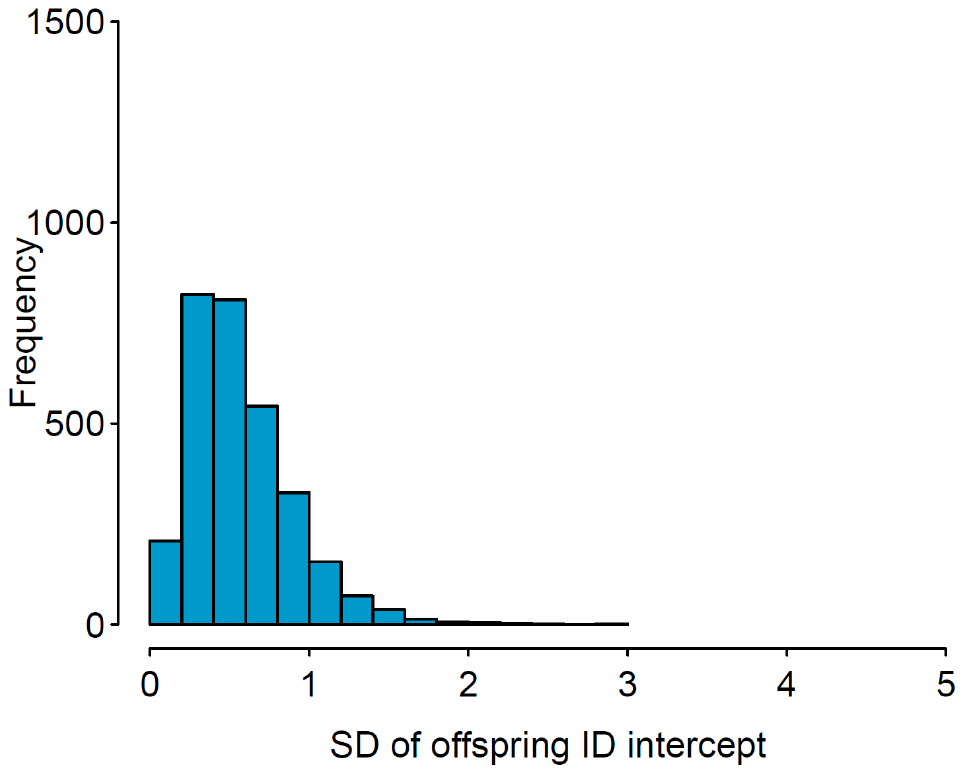 | (b) *Close proximity termination*  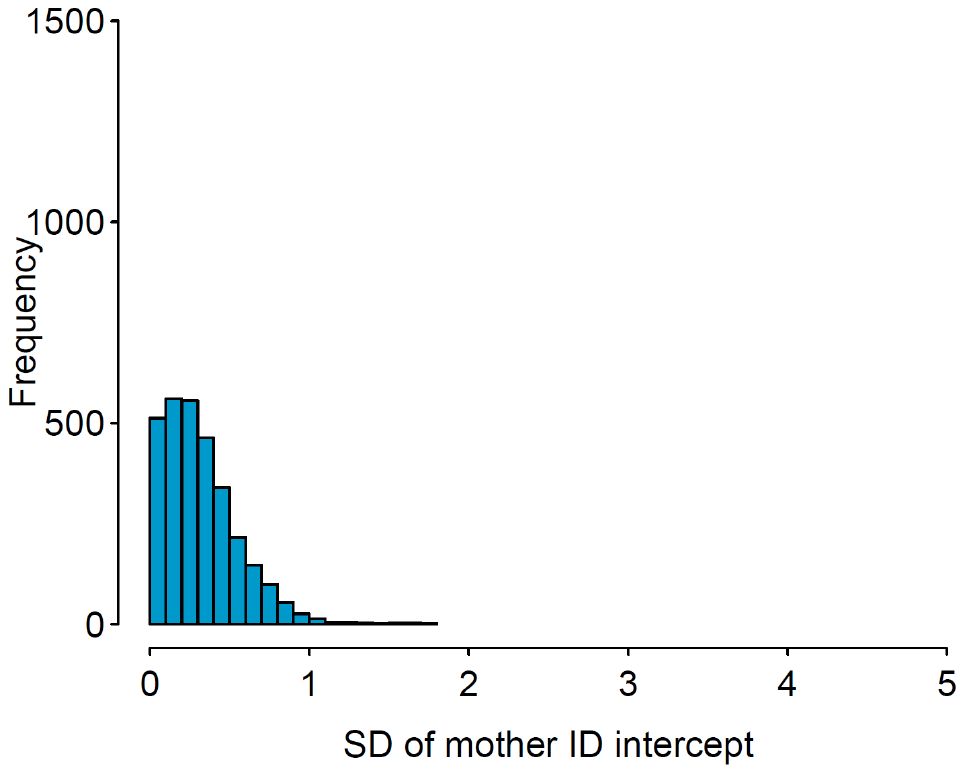  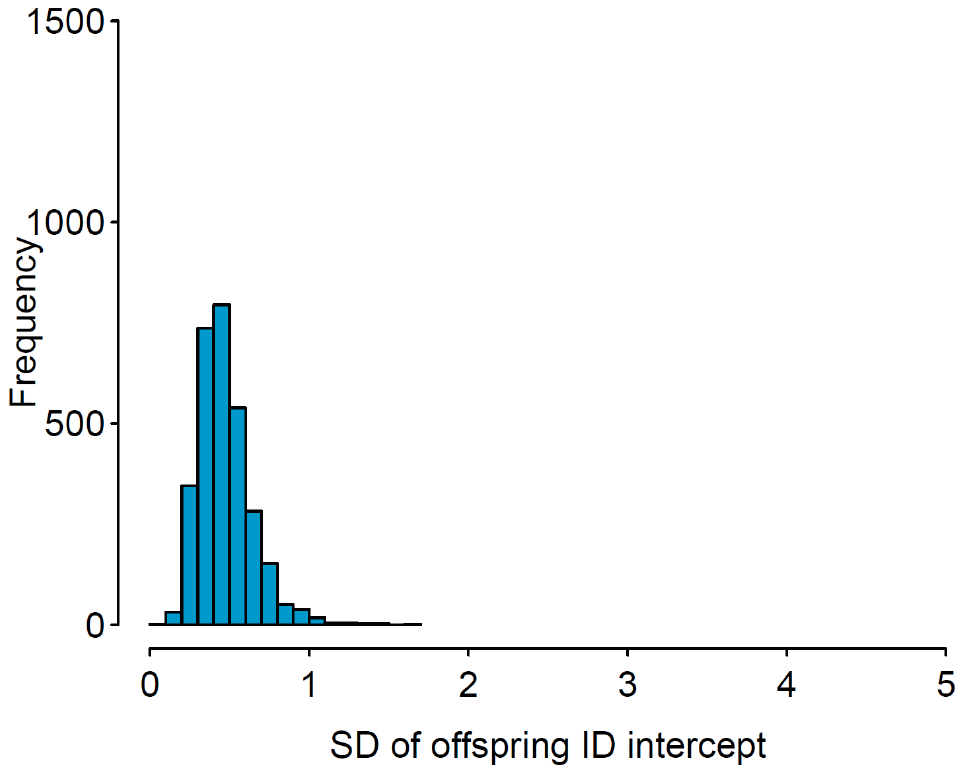 |
| --- | --- |
| (c) *Carrying*  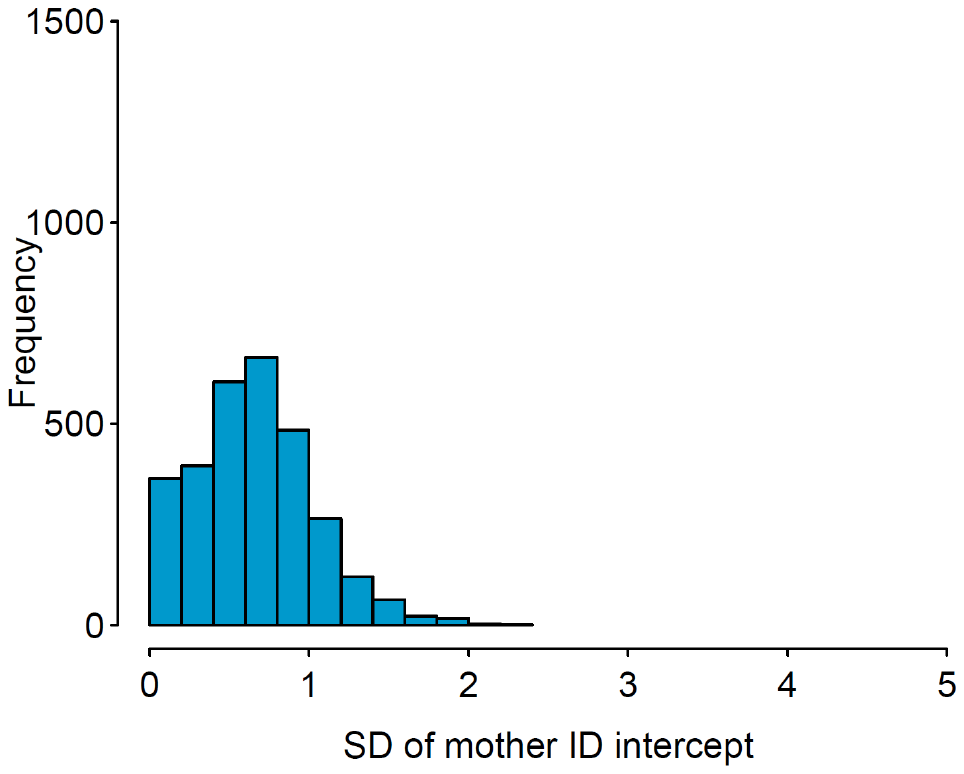  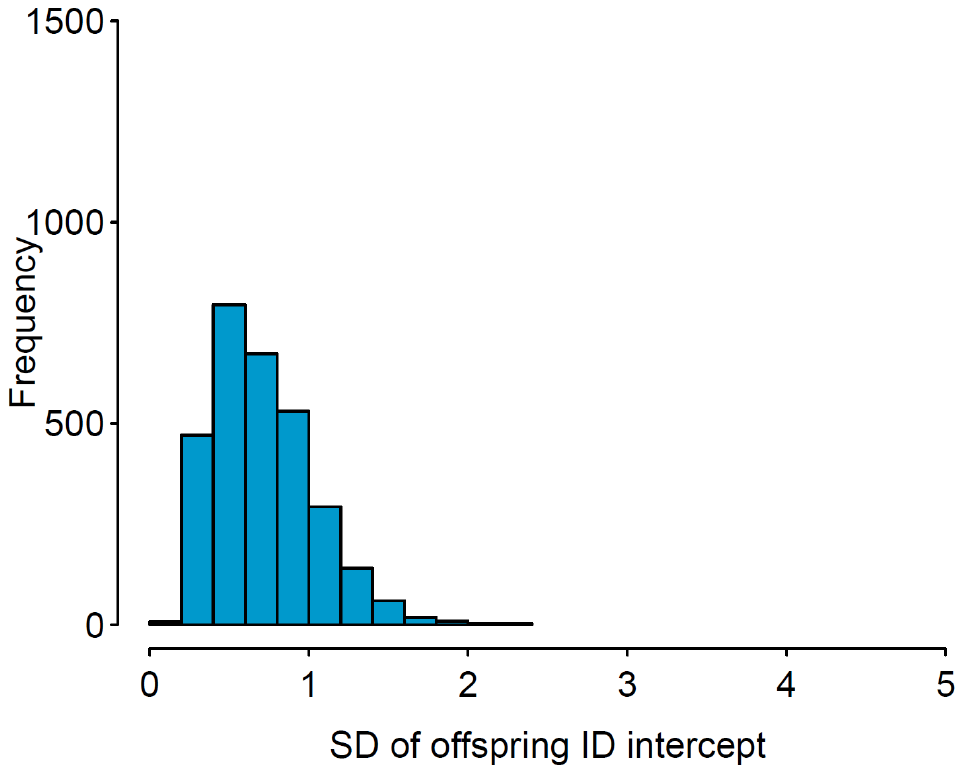 | (d) *Feeding in close proximity*  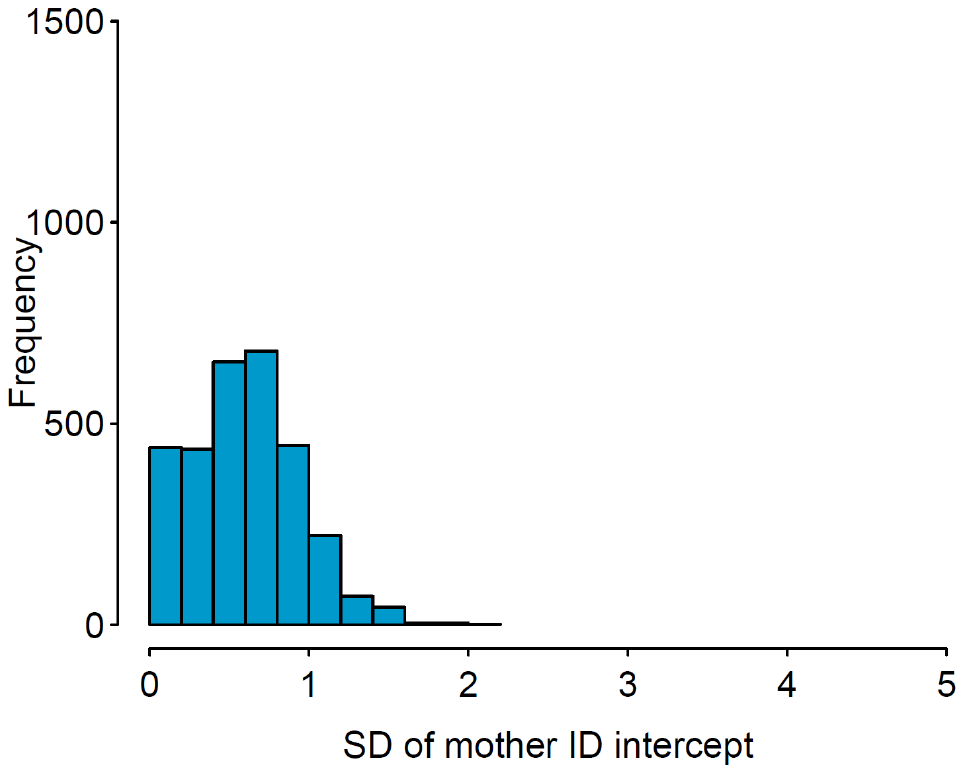  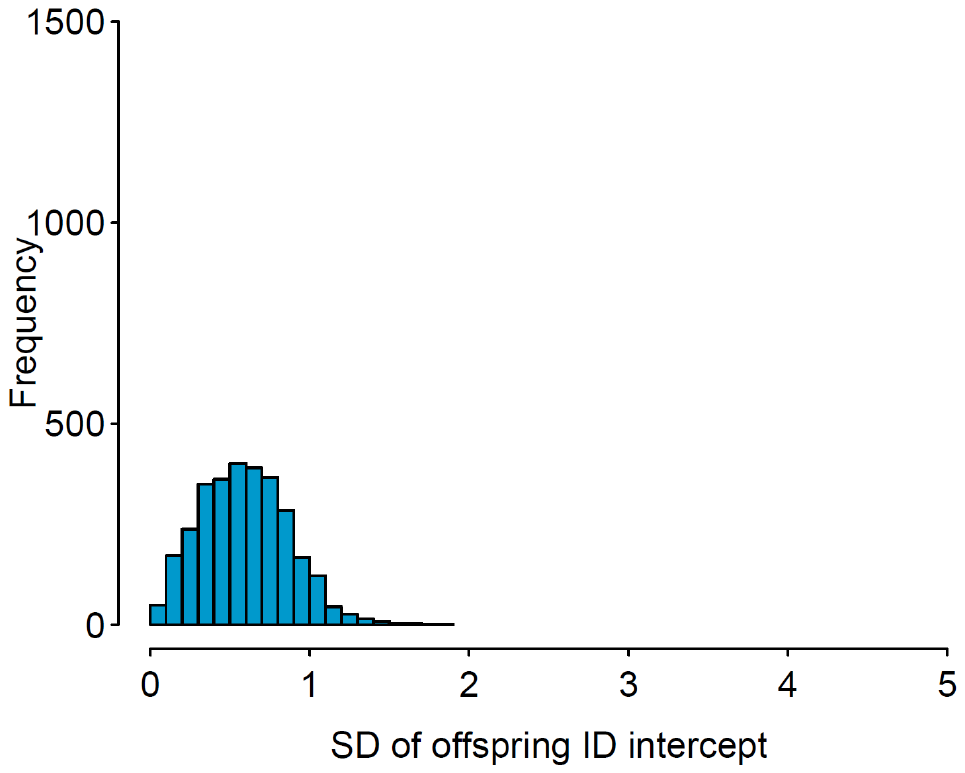 |

**Supplementary Material 6, Figure S1.** Distribution of posterior draws of the standard deviation of the random intercept of mother identity and offspring identity for the behaviours (a) contact termination (random intercept model), (b) close proximity termination (full model), (c) carrying (full model), and (d) feeding in close proximity (full model).

**Supplementary Material 7**

*Similarity in the average expression of maternal behaviour across behaviours for the individual mothers*

| **BCT** | **BCT** |  |  |
| --- | --- | --- | --- |
| **CPT** | 0.46 (0.16) | **CPT** |  |
| **CAR** | -0.43 (0.18) | -0.16 (0.64) | **CAR** |
| **FCP** | 0.01 (0.98) | -0.51 (0.11) | -0.64 (**0.03**) |

**Supplementary Material 7, Table S1.** Correlation (*r*) matrix between the average expression of behaviour (i.e., individual random intercept estimates) for all combinations of the behaviours in which the addition of mother identity improved model predictions and the corresponding *P* values. *P* value in bold denotes significant correlation. Random intercept estimates of only those mothers who were present in all the four behaviours were correlated (*N*=11 mothers). Estimates were obtained from the more parsimonious model for each behaviour. BCT: body contact termination; CPT: close proximity termination; CAR: carrying; FCP: feeding in close proximity.

**Supplementary Material 8**

*Posterior distribution of the estimated standard deviation for the random slope of offspring age within random intercept of mother identity and offspring identity nested within mother identity*

| (a) *Close proximity termination*  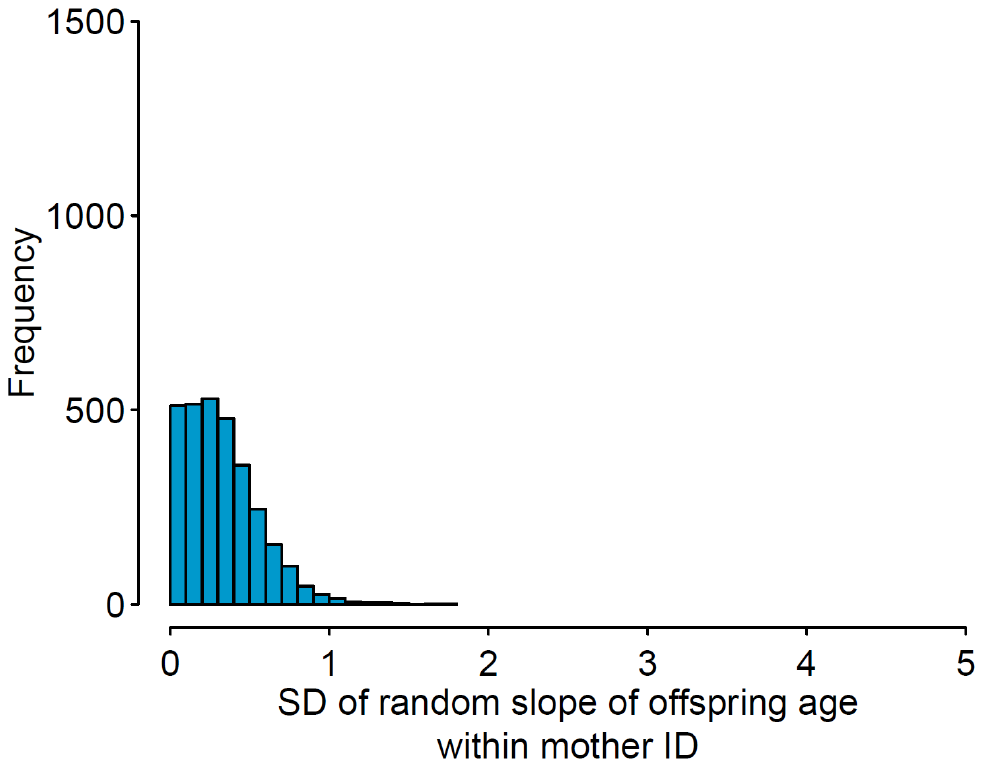  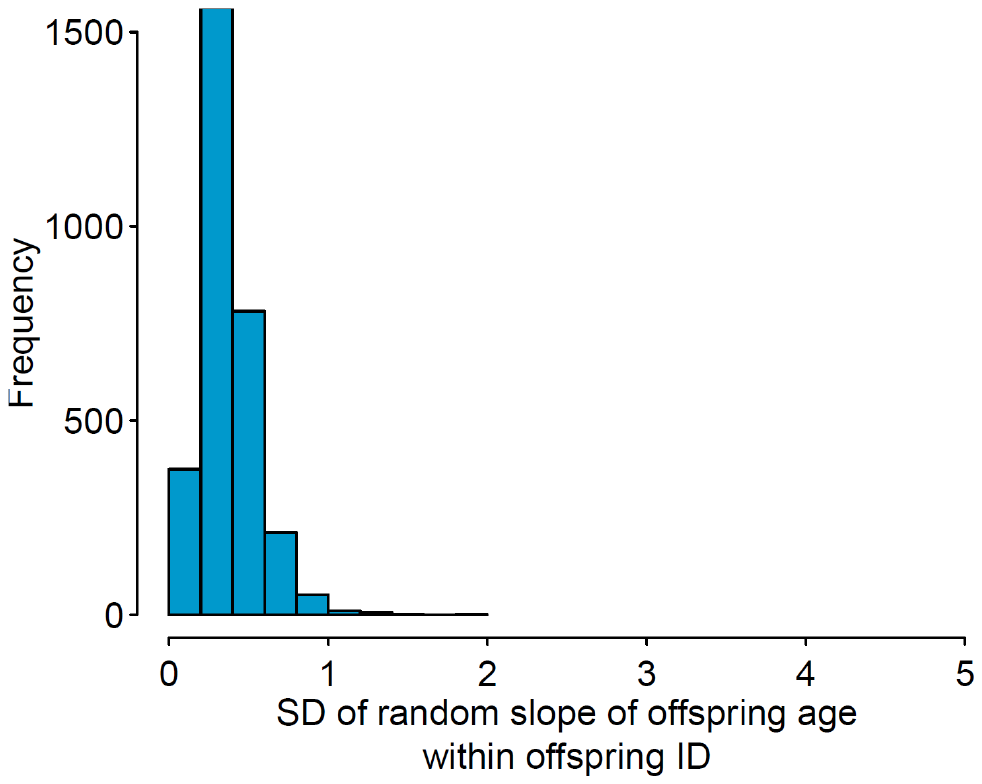 | (b) *Carrying*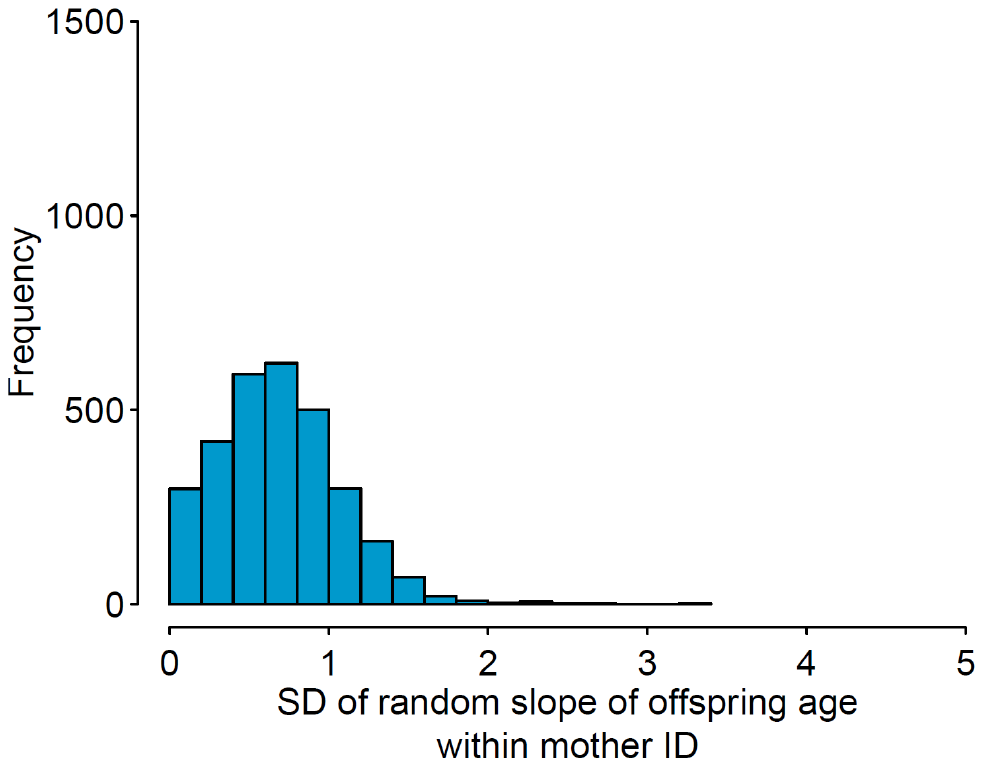  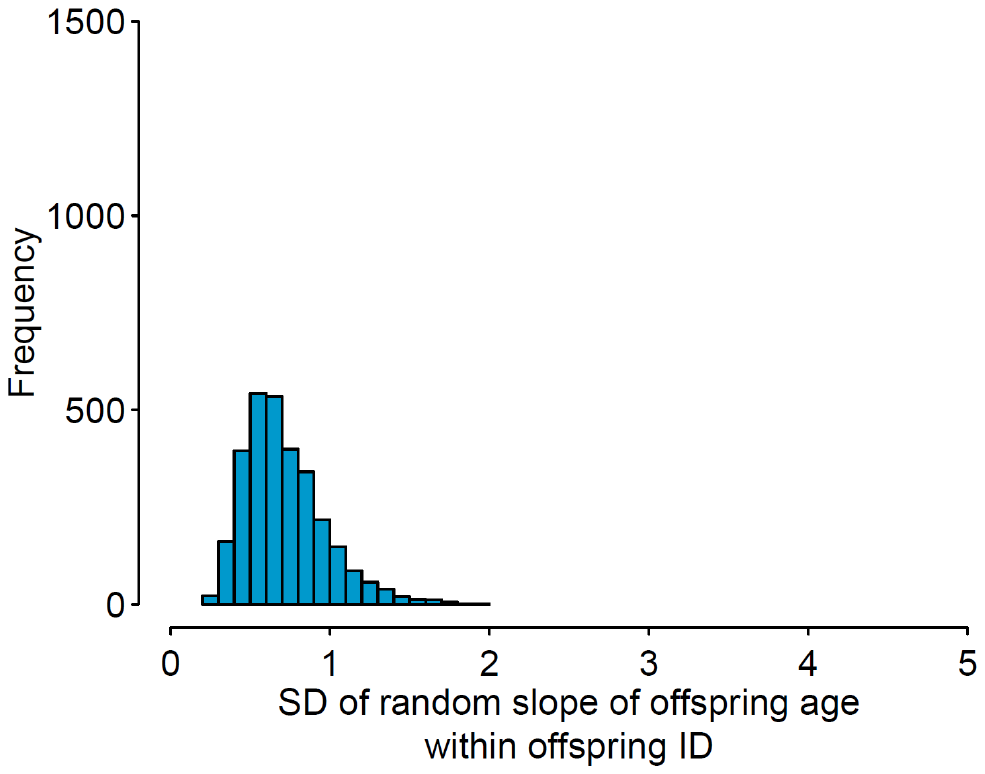 |
| --- | --- |
| (c) *Feeding in close proximity*  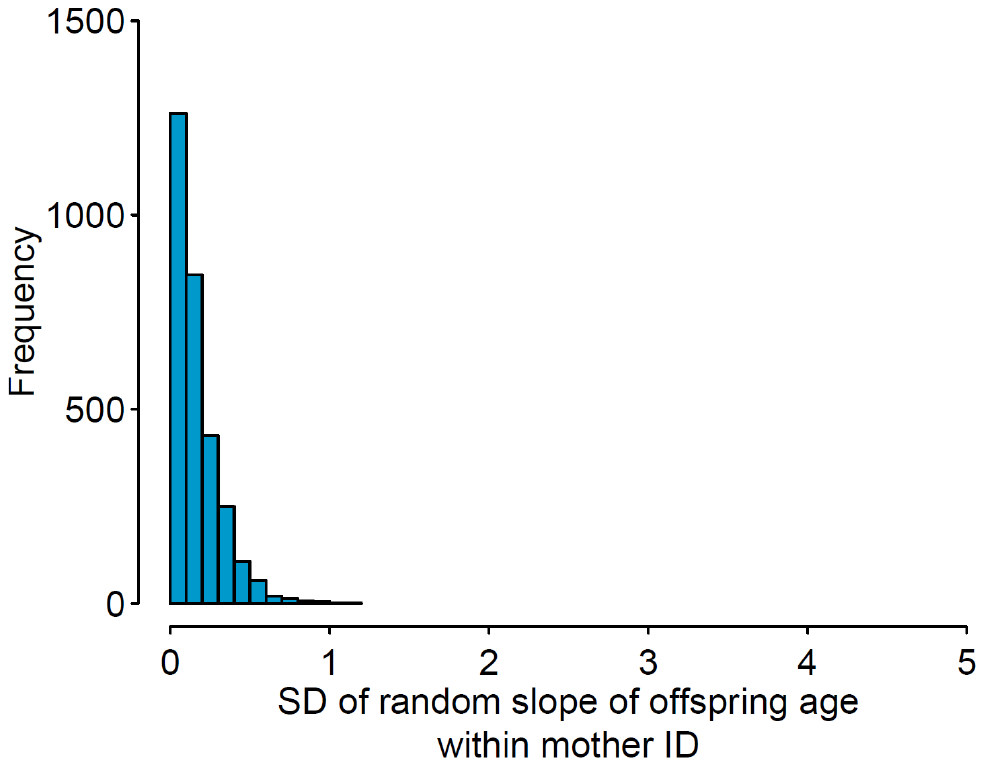  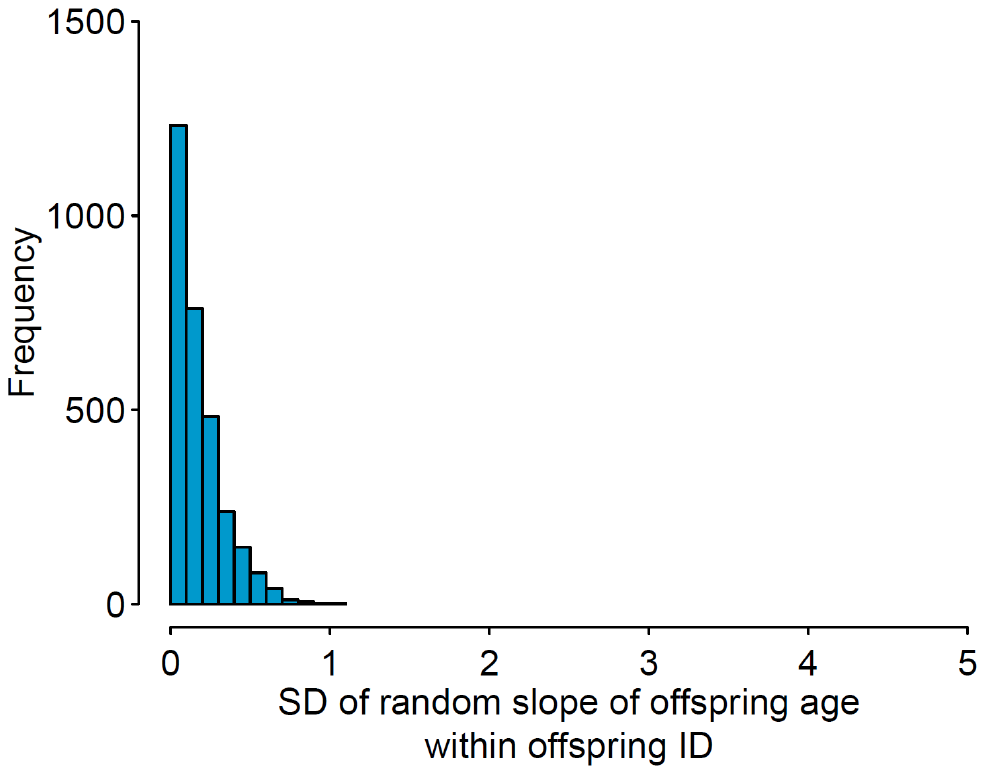 |  |

**Supplementary Material 8, Figure S1.** Distribution of the posterior draws of the standard deviation of the random slope of offspring age within mother identity and offspring identity for the behaviours (a) close proximity termination, (b) carrying, and (c) feeding in close proximity from the respective full model.

**Supplementary Material 9**

*Results of the GLMM performed using the filtered data*

**Supplementary Material 9, Table S1**. Difference in predictive performance of the fitted models using filtered data (i.e., only mothers with multiple offspring). Results of the model comparisons based on LOO-CV are shown. The most parsimonious model is marked with an asterisk (*). Δ*ELPD* ± 2 × Δ*SE* for each model was obtained by comparing it with the full model.

| Model | *ELPD* | *SE* | Δ*ELPD* | Δ*SE* | Δ*ELPD* ± 2 × Δ*SE* |
| --- | --- | --- | --- | --- | --- |
| ***Carry*** | | | | | |
| Full model* | 1472.8 | 40.7 | **0** | **0** | **-** |
| Random intercept model | 1440.8 | 40.3 | -32 | 7.2 | [-46.4, -17.6] |
| Fixed effects model | 1408.6 | 40.7 | -64.2 | 11.1 | [-86.4, -42] |
| ***Feeding in close proximity*** | | | | | |
| Full model* | 985.1 | 52.2 | **0** | **0** | **-** |
| Random intercept model | 958.5 | 50.6 | -26.6 | 8 | [-42.6, -10.6] |
| Fixed effects model | 922.8 | 49.9 | -62.3 | 12.7 | [-87.7, -36.9] |

**Supplementary Material 9, Table S2.** Estimated between-individual variation in the average expression of a behaviour and between-individual variation in behavioural plasticity for the models fitted with filtered data (i.e., only mothers with multiple offspring). Means and standard deviations of the posterior distribution of parameters, their 95% credible intervals (CI), along with ESS for the mother and offspring random effects are shown. MotherID:OffspringID denotes nested random effect.

| Random effect | Estimate | Estimated  Error | 95% CI | Rhat | Bulk  ESS | Tail  ESS |
| --- | --- | --- | --- | --- | --- | --- |
| *Carrying*; Full model; *N*=583 focal follows; *R*^2^=0.85 | | | |  |  |  |
| ~MotherID (Number of levels: 6) | | |  |  |  |  |
| SD_(Intercept)_ | 0.4 | 0.33 | [0.02, 1.23] | 1.00 | 1120 | 1579 |
| SD_(Offspring Age)_ | 0.44 | 0.4 | [0.01, 1.51] | 1.00 | 1230 | 1431 |
| Cor_(Intercept,Offspring Age)_ | -0.18 | 0.58 | [-0.97, 0.91] | 1.00 | 2360 | 2149 |
| ~MotherID:OffspringID (Number of levels: 13) | | | |  |  |  |
| SD_(Intercept)_ | 0.53 | 0.24 | [0.21, 1.12] | 1.00 | 1110 | 1660 |
| SD_(Offspring Age)_ | 0.84 | 0.28 | [0.44, 1.49] | 1.00 | 1303 | 1971 |
| Cor_(Intercept,Offspring Age)_ | -0.76 | 0.26 | [-1.0, -0.01] | 1.00 | 1293 | 1721 |
| *Feeding in close proximity*; Full model; *N*=602 focal follows; *R*^2^=0.37 | | | | |  |  |
| ~MotherID (Number of levels: 6) | | |  |  |  |  |
| SD_(Intercept)_ | 0.36 | 0.33 | [0.01, 1.24] | 1.00 | 1082 | 955 |
| SD_(Offspring Age)_ | 0.28 | 0.27 | [0.01, 0.97] | 1.00 | 1227 | 1514 |
| SD_(Offspring Age^2)_ | 0.32 | 0.29 | [0.01, 1.06] | 1.00 | 1339 | 1867 |
| Cor_(Intercept,Offspring Age)_ | 0.02 | 0.5 | [-0.87, 0.88] | 1.00 | 2684 | 2233 |
| Cor_(Intercept,Offspring Age^2)_ | -0.07 | 0.5 | [-0.9, 0.86] | 1.00 | 2673 | 2103 |
| Cor_(Offspring Age,Offspring Age^2)_ | -0.01 | 0.49 | [-0.87, 0.87] | 1.00 | 2588 | 2465 |
| ~MotherID:OffspringID (Number of levels: 13) | | | |  |  |  |
| SD_(Intercept)_ | 0.45 | 0.23 | [0.07, 1.02] | 1.00 | 1060 | 1007 |
| SD_(Offspring Age)_ | 0.22 | 0.21 | [0.01, 0.8] | 1.00 | 1187 | 1533 |
| SD_(Offspring Age^2)_ | 0.7 | 0.25 | [0.33, 1.3] | 1.00 | 1072 | 1661 |
| Cor_(Intercept,Offspring Age)_ | 0.09 | 0.51 | [-0.85, 0.92] | 1.00 | 2551 | 2098 |
| Cor_(Intercept,Offspring Age^2)_ | 0 | 0.41 | [-0.73, 0.78] | 1.00 | 884 | 1208 |
| Cor_(Offspring Age,Offspring Age^2)_ | 0.07 | 0.5 | [-0.85, 0.91] | 1.01 | 513 | 1223 |

| (a) *Carrying*  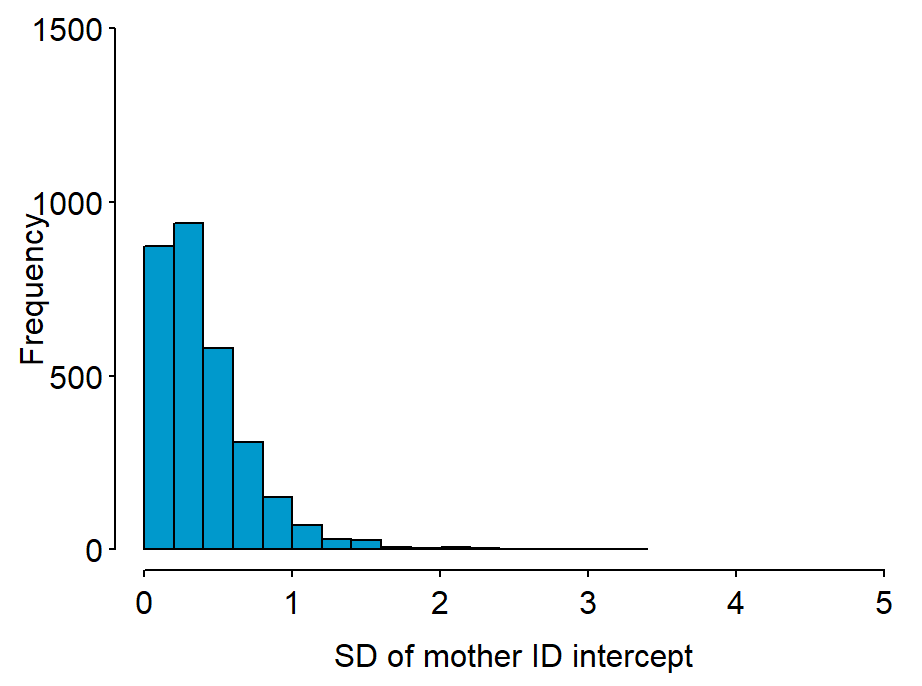  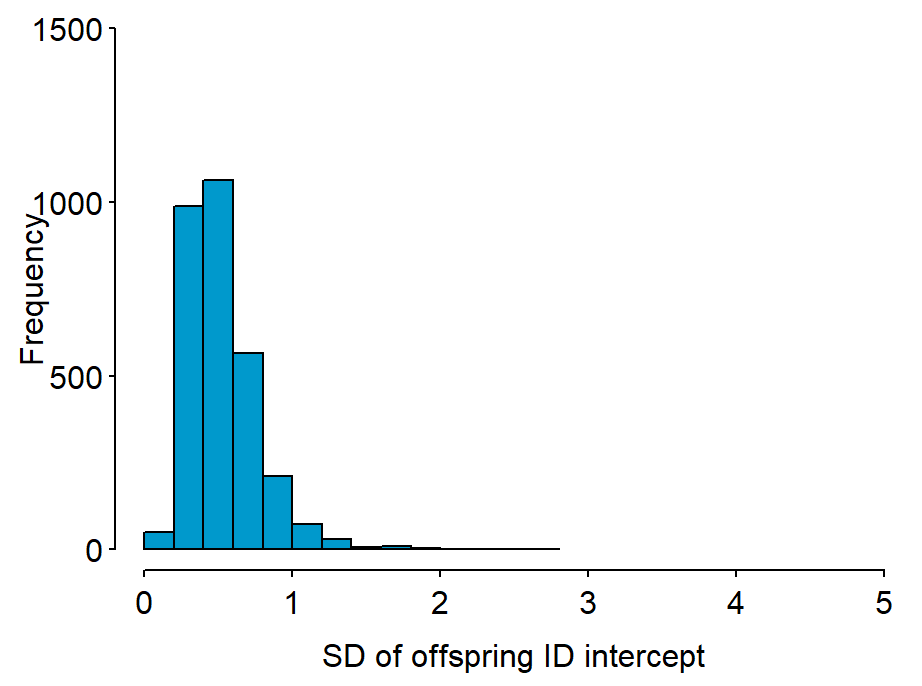 | (b) *Feeding in* close *proximity*  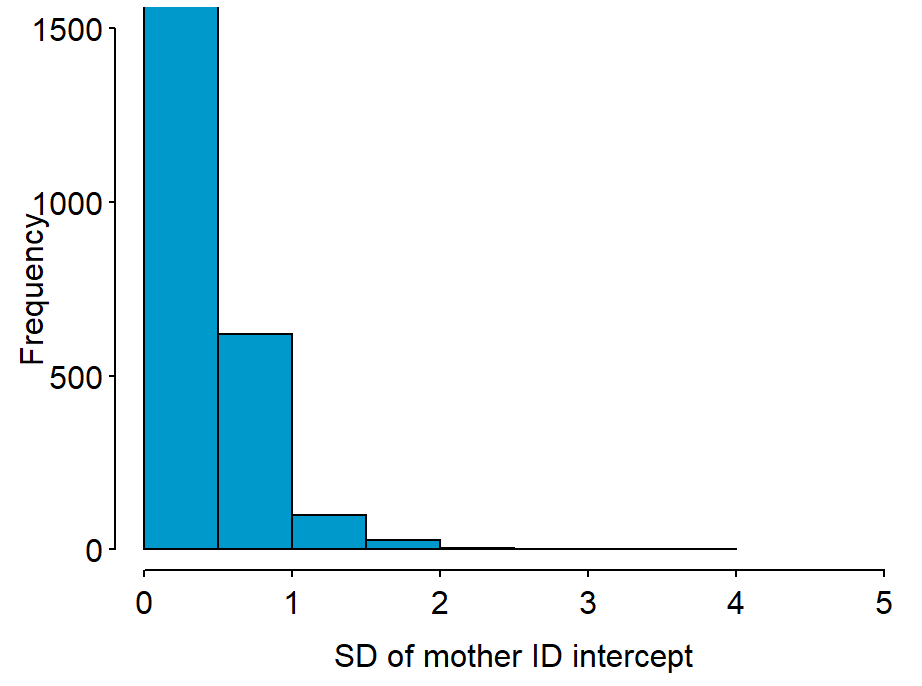  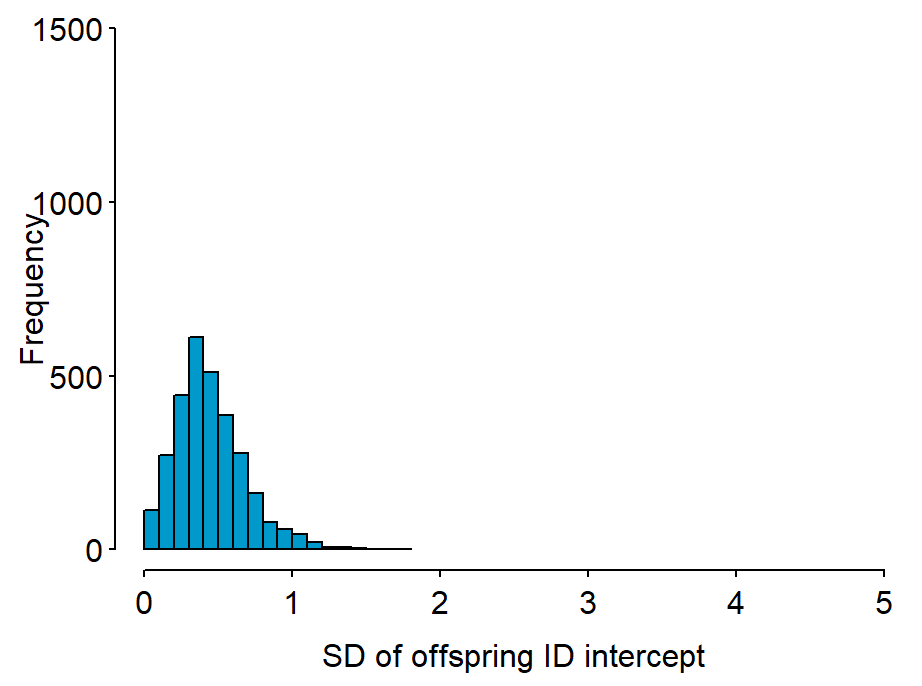 |
| --- | --- |

**Supplementary Material 9, Figure S1.** Distribution of posterior draws of the standard deviation of the random intercept of mother identity and offspring identity for the behaviours (a) carrying and (b) feeding in close proximity from the respective full model fitted with filtered data (i.e., only mothers with multiple offspring).

| (a) *Carrying*  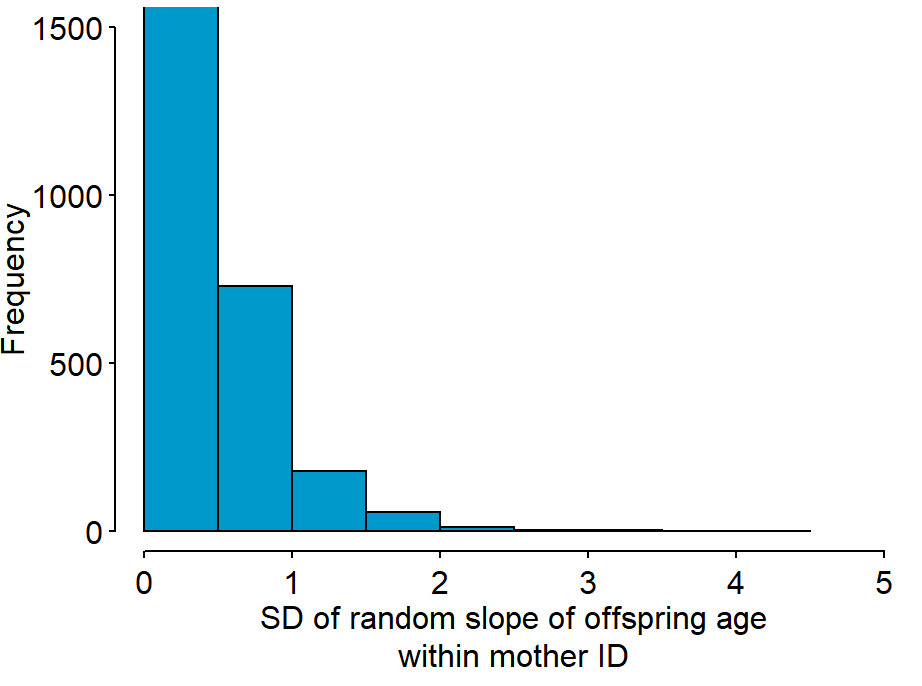  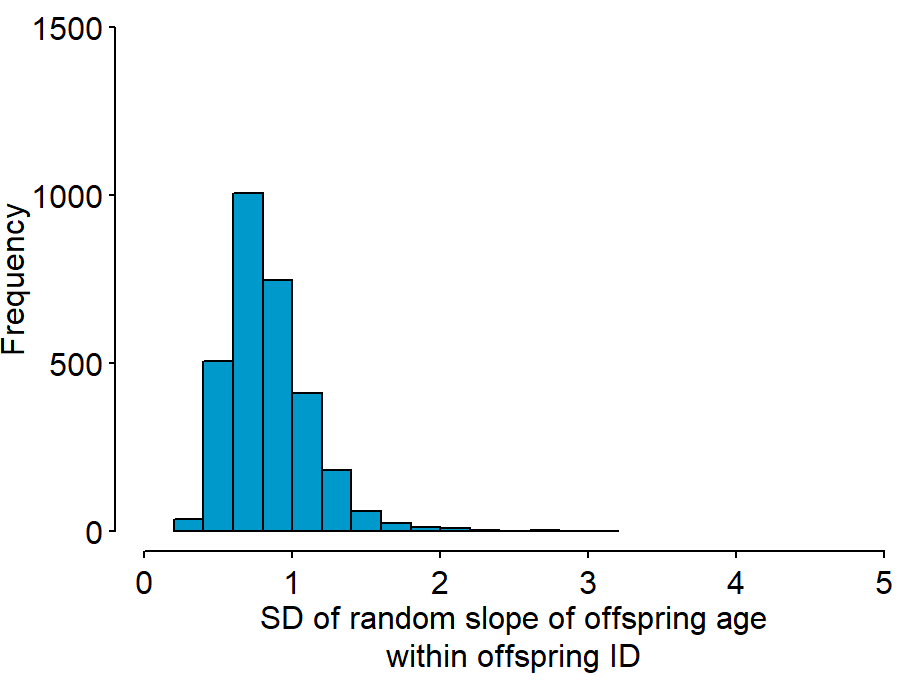 | (b) *Feeding in* close *proximity*  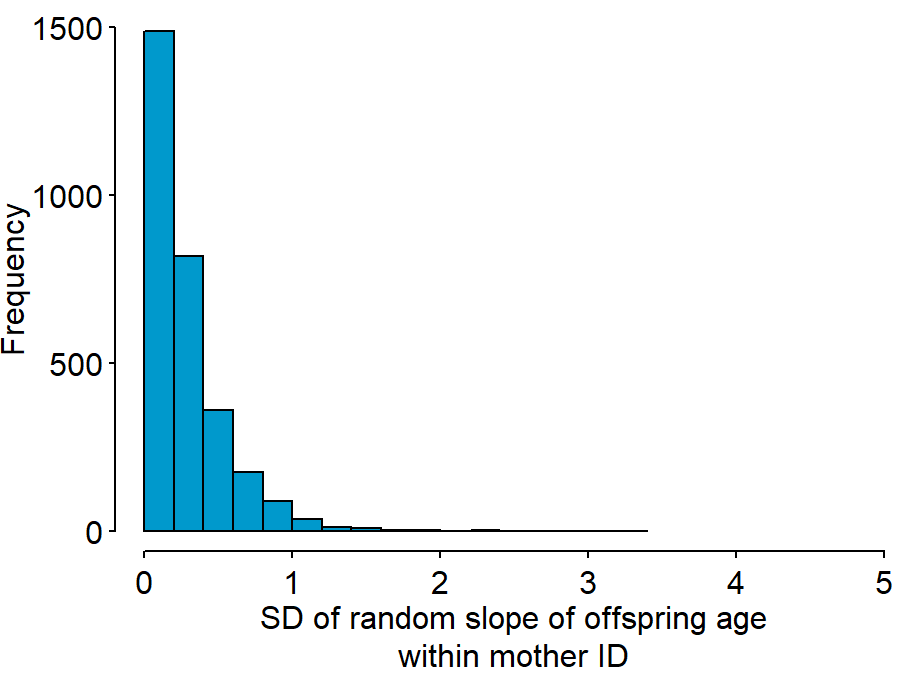  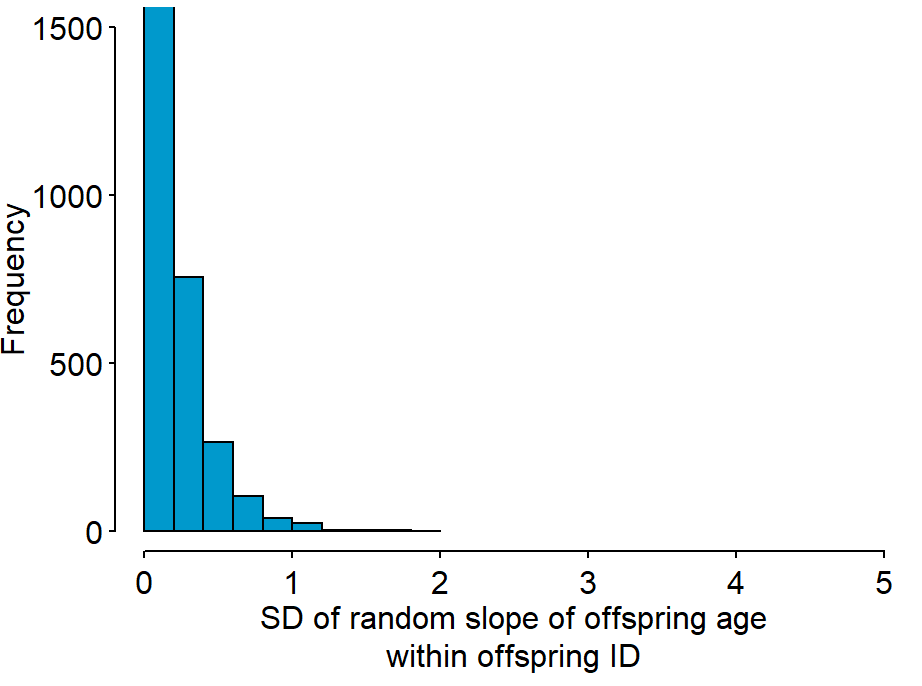 |
| --- | --- |

**Supplementary Material 9, Figure S2.** Distribution of the posterior draws of the standard deviation of the random slope of offspring age within mother identity and offspring identity for the behaviours (a) carrying and (b) feeding in close proximity from the respective full model fitted with filtered data (i.e., only mothers with multiple offspring).
